# Single-Cell eQTL Mapping Reveals Cell Type-Specific Genetic Regulation in Lung Cancer

**DOI:** 10.1101/2025.04.10.646767

**Authors:** Yating Fu, Yi Wang, Chang Zhang, Chen Jin, Jiaying Cai, Linnan Gong, Chenying Jin, Chen Ji, Yuanlin Mou, Caochen Zhang, Shihao Wu, Xinyuan Ge, Yahui Dai, Sunan Miao, Huimin Ma, Xiaoyang Ma, Mengping Wang, Lijun Bian, Erbao Zhang, Juncheng Dai, Zhibin Hu, Guangfu Jin, Meng Zhu, Hongbing Shen, Hongxia Ma

## Abstract

Genome-wide association studies (GWAS) have identified over 50 lung cancer risk loci; however, the precise cellular context of these genetic mechanisms remains unclear due to limitations in bulk tissue eQTL analyses. Here, we present the largest single-cell eQTL atlas of human lung tissue to date, profiling 222 donors using multiplexed scRNA-seq. We identified 16,785 eQTLs across 17 cell types, with over 90% of sc-eQTLs and 59% eGene being cell-type-specific, and fewer than 23% were detectable in paired bulk datasets. Integration with GWAS for non-small cell lung cancer highlighted epithelial and immune cells as key contributors to genetic susceptibility, identifying 28 candidate genes within known risk loci and 24 in novel regions. Notably, 47% of established NSCLC susceptibility loci exhibited cell-type-specific pleiotropic genetic regulation. This study provides a valuable resource of lung sc-eQTLs and illuminates how genetic variation modulates gene expression in a cell-type-specific fashion, contributing to lung cancer susceptibility.

## Introduction

Lung cancer remains the most prevalent malignancy worldwide and the leading cause of cancer-related deaths^1^, with non-small cell lung cancer (NSCLC) accounting for approximately 85% of all cases^2^. While tobacco smoking remains the primary risk factor, genetic predisposition also plays a critical role in lung cancer development. Twin studies estimate the heritability of lung cancer at approximately 18%-26%^3,4^, highlighting the significant contribution of inherited genetic variation. Over the past two decades, genome-wide association studies (GWAS) have identified over 50 susceptibility loci for NSCLC across diverse populations^5–13^, implicating candidate genes involved in smoking behavior (e.g., *CYP2A6*, *CHRNA5*, *CHRNB4*), DNA repair pathways (e.g., *CHEK2*, *BRCA2*, *ATM*), and telomere maintenance (e.g., *TERT*, *RTEL*, *OBFC1*)^14^. Nevertheless, the functional mechanisms and causal genes underlying most GWAS loci remain poorly characterized, underscoring a critical gap in our understanding of NSCLC susceptibility.

Identifying target genes from GWAS loci remains a major challenge, primarily because most susceptibility variants reside in non-coding genomic regions, particularly within regulatory elements^15^. These variants typically regulate gene expression rather than directly altering protein-coding sequences^16^. To address this, expression quantitative trait loci (eQTL) analyses have emerged as a primary strategy to link GWAS variants to their putative target genes^17^. However, most prior studies have relied on bulk RNA sequencing, which captures gene expression as an average across heterogeneous cell populations. This averaging effect masks cell-type-specific regulatory mechanisms, particularly when variants exert their effects in distinct cellular contexts or display divergent regulatory activities across different cell types.^18–20^.

Recent advances in single-cell RNA sequencing (scRNA-seq) offers an unbiased approach to characterize cellular composition and cell-type-specific gene expression, providing unprecedented resolution into intra-individual cellular heterogeneity. Emerging evidence demonstrates that single-cell eQTLs (sc-eQTLs) can uncover cell-type-specific regulatory variants that are often missed in bulk eQTL analyses^21–25^. Despite several sc-eQTL databases have been developed, few studies have focused specifically on lung tissues^26,27^. The lung consists of diverse cell types, including alveolar type II (AT2), AT1, club, basal, and neuroendocrine cells—key progenitor cells in lung cancer—which interact dynamically with microenvironmental components such as macrophages, ciliated cells, and immune cells during tumorigenesis^28–31^. A recent study by Long et al. integrated single-nucleus RNA-seq (snRNA-seq) and single-nucleus ATAC-seq (snATAC-seq) to investigate genetic regulation underlying lung cancer susceptibility^32^. However, their indirect identification of target genes highlights the critical need for sc-eQTL mapping to elucidate the mechanisms of lung cancer susceptibility. Although Natri et al. had mapped sc-eQTLs in 114 lung samples, 58% were derived from pulmonary fibrosis patients, limiting generalizability to normal lung regulatory networks^21^.

Here, we aimed to identify target genes associated with NSCLC at the cellular subpopulation level. Using a pooled multiplexing strategy, we leveraged scRNA-seq data from 222 adult lung samples from Chinese individuals to construct the first sc-eQTLs atlas for normal lung tissues. This resource is publicly available at http://ccra.njmu.edu.cn/LungSCeQTL. We then integrated cell-type-specific eQTLs with a Chinese NSCLC GWAS (14,240 NSCLC cases and 14,813 controls) to pinpoint target genes at GWAS loci across various cell types. Finally, we identified novel NSCLC susceptibility genes through integrative biological analyses and functional assays.

## Results

### The Characteristic of Chinese LungSCeQTL cohort

We investigated the cell type-specific effects of genetic variants on gene expression within the Chinese Lung Single Cell eQTL (LungSCeQTL) cohort (**Figure 1A**). This cohort comprised 226 Chinese individuals who underwent lung resection surgery for either lung cancer (n=188) or non-neoplastic lung diseases (e.g., pulmonary bullae; n=38) between October 2021 to July 2022. Paired blood samples and lesions-distant normal lung tissues were collected from all participants.

**Figure 1.**
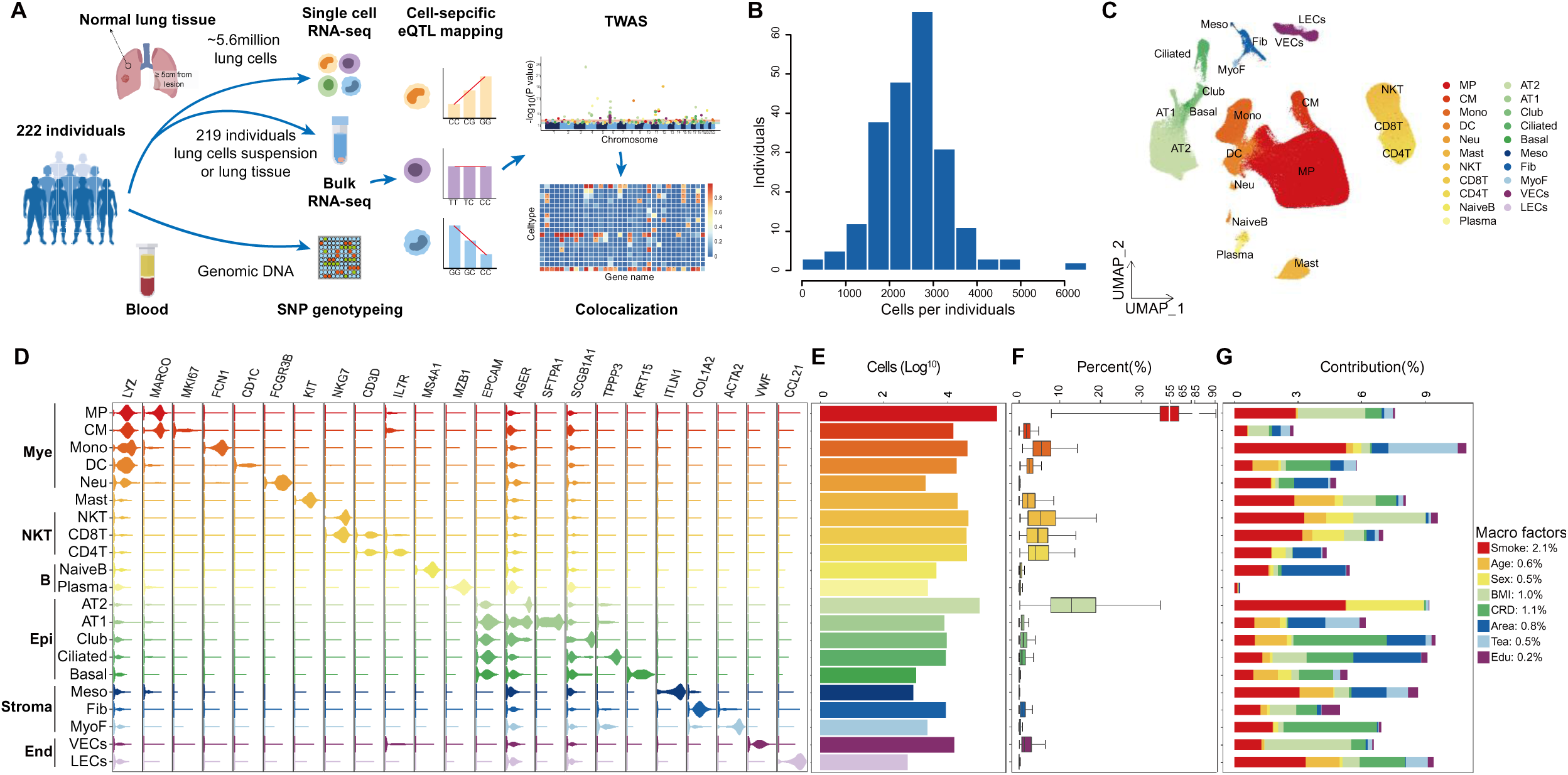
Human lung cell atlas in the Chinese LungSCeQTL cohort. **(A)** Study design overview. **(B)** The mean of 2,533 individual cells per donor, ranged from 330 to 10,184 after scRNA-seq, demultiplexing, and quality-control filtering. **(C)** UMAP of 562,255 cells across 222 individuals, with 21 transcriptionally distinct populations. **(D)** Violin plot visualizing the normalized RNA expression of selected canonical marker genes across cell types. **(E)** Total number of cells per cell type after sequencing, demultiplexing, and quality-control filtering. (**F)** Total percentage of each cell type as a proportion of the overall population for each individual. **(G)** Contributions of the macro-variables to the effect on the cellular composition of lung tissue.

Genotyping was performed using the Infinium Asian Screening Array (ASA), yielding 657,490 single nucleotide polymorphisms (SNPs). After quality control and imputation against the 1000 Genomes Project reference panel, 4,838,930 SNPs with an INFO > 0.8 and minor allele frequency (MAF) > 0.01 were retained for further analysis. Fresh normal lung tissues were dissociated into single-cell suspensions, and scRNA-seq data from 870,061 cells were generated using a pooled multiplexing strategy^33^ across 29 experimental pools. To assess batch effects across different pools, repeated lung tissues from six patients were sequenced in separate pools.

After demultiplexing, removal of doublets, and quality control, 562,255 cells from 222 individuals were retained, with an average of 2,533 cells per donor (range: 330-10,184; **Figure 1B**). All individuals were unrelated (identity-by-descent [IBD] < 0.125), with a mean age of 55.67 years, 57.7% female, and 70.7% non-smokers.

### Lung Single Cell Atlas in the Chinese LungSCeQTL cohort

Data integration, dimensionality reduction, and unsupervised clustering of qualified cells were performed using the Seurat package (**Methods**)^34^. Based on canonical marker gene expression for lung tissue cell types from previous studies ^31,35^, cells were classified into seven major lineages. The cell compositions ranged from 0.99% (B cells) to 60.52% (myeloid cells). Reanalysis within each pool revealed consistent distributions and similar cell proportions across pools. Notably, gene expression profiles for the seven lineages exhibited a high correlation (R > 0.8) between the six replicated samples, confirming the stability of our pooled multiplexing strategy with minimal batch effects.

We further refined these major lineages into 21 distinct cell types using a combination of hierarchical supervised and unsupervised classification methods. Visualization of cell types using Uniform Manifold Approximation and Projection (UMAP) revealed the hierarchical relationships among these cell types (**Figure 1C**). Cell-type-specific marker genes derived from previous studies showed specific high expression in the corresponding cell types (**Figure 1D**). Among the 21 identified cell types, macrophage cells (MP) dominated the cellular landscape (median 44.32%, IQR 34.66-56.93%), followed by AT2 (12.88%, IQR 7.83-18.84%) (**Figure 1E and F**). The range of cell type proportions were consistent with previous reports on lung tissue^36–40^.

While cell type proportions remained consistent across pools, significant inter-individual variation was observed. To systematically investigate epidemiological determinants of cellular heterogeneity, we applied the mixed-effects modeling of associations of single cells (MASC)^41^, analyzing 20 covariates previously implicated in lung cancer^42–44^. Independent cluster association p-values were aggregated into a gamma-distributed test statistic **(Methods)**, identifying eight factors significantly associated with lung tissue cell composition. Using the “Relaimpo” R package^45^, we quantified the effect size of these factors and found that smoking emerged as the predominant modulator, explaining 2.1% of cross-sample variability, with the strongest effects on monocytes (5.28%) and AT2 cells (5.24%) (**Figure 1G**).

### Mapping eQTL across cell types in the Chinese LungSCeQTL cohort

To investigate the impact of genetic variations on cell type-specific gene expression, we performed cis-eQTL analysis using FastQTL^46^ in 17 well-represented cell types, each with at least 90 donors and five or more detected cells. To account for potential confounders, we included eight variables affecting lung tissue cell composition, the first ten principal components (PCs), and the top Probabilistic Estimation of Expression Residuals (PEER) factors as covariates **(Methods)**. We conducted conditional analysis by iteratively incorporating the lead cis-eQTL SNP (eSNP) as a covariate, identifying a total of 16,785 independent eQTLs and 14,906 eGenes across the 17 cell types. Of these, 14,906 eSNPs were detected in the first round, and additional 1,879 eSNPs were identified in the second round of analysis (**Figure 2A**). The number of independent eQTLs and eGenes varied across cell types, with 5,047 eQTLs (linked to 4,145 eGenes) in MP cells and 79 eQTLs (corresponding to 79 eGenes) in MyoF cells. This variation is likely due to differences in cell capture, rather than overall sample size (**Figure 2B**), underscoring the importance of enriching for rare cell types in sc-eQTL studies.

**Figure 2.**
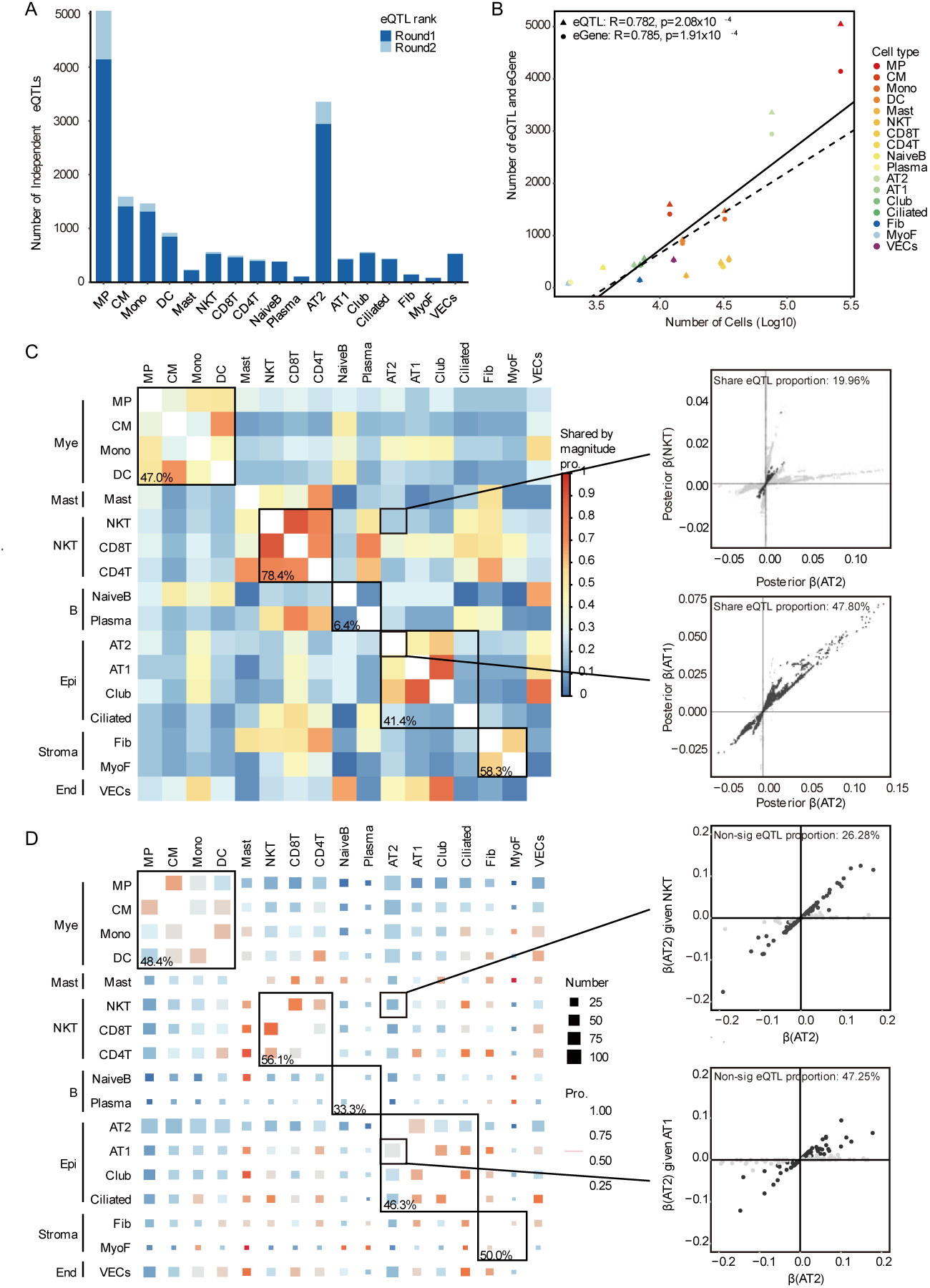
Mapping eQTL across cell types in the Chinese LungSCeQTL cohort. (**A**) The number of eSNP1 (the top SNP) and eSNP2 (the second most significant SNP) identified per cell type, shown across up to two rounds of conditional eQTL analysis. Dark blue represents eSNP1 and light blue reprents eSNP2. **(B)** Correlation between the number of independent eQTLs (triangles and solid lines) and eGenes (circles and dashed lines) per cell type with the number of cells of that cell type (Pearson’s correlation). Cell types are colored by sublineage. An eGene is defined as a gene that has an eQTL. **(C)** Pairwise sharing by the magnitude of eQTLs among cell types. For each pair of cell types, we considered the eQTLs that were significant (1fsr < 0.05) in at least one of the two cell types and plotted the proportion of these shared in magnitude (i.e., effect estimates with the same direction and within a factor of 2 in size). Brackets around cell type labels highlight cell types that show higher sharing within their respective lineage. Mean of pairwise percentage sharing per lineage is shown in black. Examples include the allelic effects of eSNPs in AT2 cells versus NKT cells (top) AT1 cells (bottom). **(D)** Heatmap shows pairwise correlations in allelic effects across cell types, indicating higher sharing among related cell lineages. The size of each square reflects the number of shared eQTL genes, and the color represents the proportion of non-significant eSNPs after conditional regression, with higher values indicating greater genetic control sharing. Examples include the allelic effects of eSNPs in AT2 cells before and after conditioning on lead eSNPs from NKT cells (top) versus AT1 cells (bottom)

Nearly 90.97% (13,114/14,416) of identified cis-eQTLs were significant in only a specific cell type. Similarly, 59.05% (5,142/8,708) of eGenes were detected exclusively in one cell type. Notably, only 6.15% (316/5,142) of these eGenes were uniquely expressed in the corresponding cell type, while the majority were expressed across an average of 16 cell types. Furthermore, we observed strong co-expression of these eGenes across related cell types, particularly within the same major lineages, conforming that most sc-eQTL-associated eGenes are driven by cell-type-specific regulatory mechanisms rather than restricted expression. For eGenes detected in only one cell type but expressed across multiple cell types, we further estimated the posterior probability of shared effects (m-value > 0.9) to eliminate potential biases arising from unbalanced sample sizes^47^. Consistently, the distribution of eGenes remained highest in a single cell type. Collectively, these findings highlight the existence of genuine regulatory heterogeneity across lung cell types.

Using multivariate adaptive shrinkage (MASHR)^48^, we evaluated the extent of shared eQTL effects across cell types. We found that a median of 94.1% (IQR: 91.6-96.1%) of eQTL signals were concordant in direction (i.e., effect estimates with the same sign), while 23.3% (IQR: 14.1-41.7%) were also shared in magnitude (i.e., effect estimates differing by less than twofold) across cell-type pairs (**Figure 2C**). We observed higher proportions of shared by magnitude within major lineages; for instance, 47.80% of eQTLs were shared between AT2 and AT1 cells, compared to only 19.96% between AT2 and NKT cells (**Figure 2C**).

Nearly 40.95% (3,566/8,708) of eGenes were identified in two or more cell types, with 85.15% of their lead eSNPs differing between cell types. To determine whether these eSNPs reflect independent regulatory effects versus linkage disequilibrium (LD)-driven associations, we performed conditional analysis by regressing lead eSNPs from one cell type against another across all 272 cell type pairs (**Methods**). The proportion of non-significant eSNPs after regression quantified shared genetic control of gene expression^22^, with higher values indicating stronger regulatory overlap (**Figure 2D**). Lineage-intrinsic pairs showed higher proportions of non-significant eQTLs, suggesting that genetic regulation is more commonly shared among closely related cell types. For instance, 47.25% (86/182) of shared eQTLs between AT2 and AT1 cells became non-significant after regressing out lead eSNPs from AT1, compared with only 26.28% (41/156) between AT2 and NKT cells (**Figure 2D**). Notably, non-significant eSNPs after regression analysis were in strong LD (r² > 0.75), while retained significant eSNPs showed weaker LD (r² < 0.25).

### Comparison of single-cell eQTLs with bulk eQTL and previous studies

A total of 219 paired lung tissues also underwent bulk RNA-seq (Bulk219) in our study. We compared the identified sc-eQTLs with those from the Bulk219 dataset. The concordance of allelic direction across all tested loci was 74.4% (ranging from 57.0% in NaiveB cells to 85.1% in MP cells) (**Figure 3A**). However, only 22.8% (ranging from 1.3% in MyoF cells to 31.4% in CM cells) of sc-eQTLs were statistically significant at FDR < 0.05 in the Bulk219 dataset (**Figure 3B**), mapping to 43.3% (ranging from 39.2% in MyoF cells to 64.5% in CM cells) of eGenes. Reverse analysis revealed that 59.6% of eGenes and 37.9% of eQTLs identified in the Bulk219 dataset were statistically significant in the sc-eQTLs at FDR < 0.05, predominantly in a single cell type (**Figure 3C and D**). Further conditional analysis revealed that adjusting for Bulk219 eSNPs rendered 34.7% of sc-eQTL associations non-significant (range: 6.3% [NaiveB] to 47.1% [MP]), while reciprocal adjustments for sc-eQTLs similarly reduced significance in Bulk219. As expected, the extent of genetic regulation shared between cell types and Bulk219 significantly correlated with the abundance of each cell type in our dataset. Collectively, these findings underscore the power of sc-eQTL mapping in identifying cell-type-specific regulatory variants that are largely undetectable in bulk eQTL analyses.

**Figure 3.**
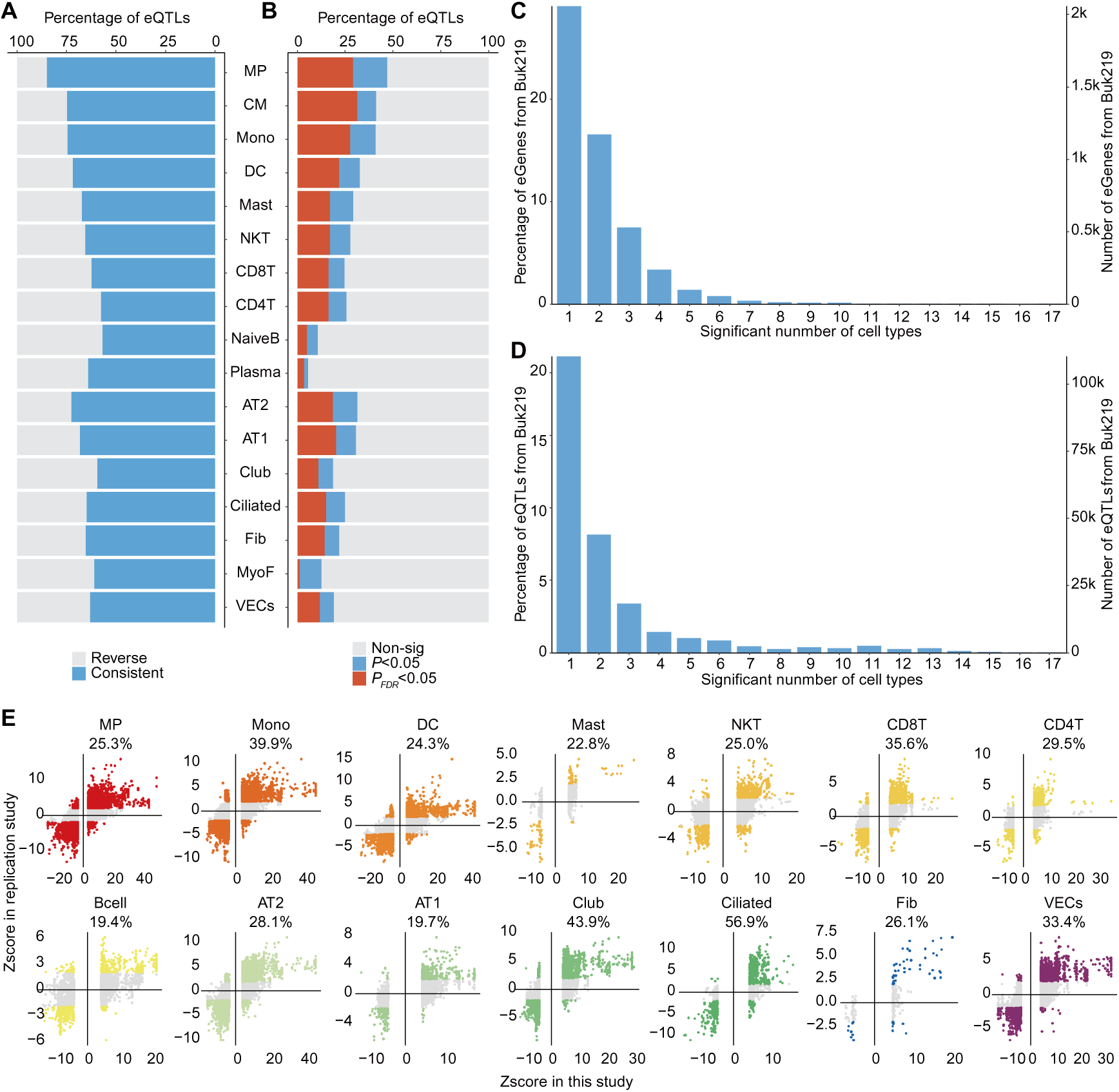
Comparison of single-cell eQTLs with bulk eQTL and previous studies. **(A)** Concordance of allelic direction of effect between independent single-cell eQTLs and the Bulk219 eQTL dataset. Blue bars indicate concordant eQTLs, while light gray bars show those with reversed effects. **(B)** Percentage of significant single-cell eQTLs replicating in the Bulk219 eQTL dataset, at an FDR threshold of 0.05 and P-value threshold of 0.05. Red bars indicate eQTLs with FDR < 0.05, while blue and light gray show those with P<0.05 and no significant, respectively. **(C)** The percentage and number of cell types with overlapping eGenes by replicating the Bulk219 cis-eQTLs in sc-eQTLs. **(D)** The percentage and number of cell types with overlapping eSNPs by replicating the Bulk219 cis-eQTLs in sc-eQTLs. **(E)** The Z-scores for eQTLs in the Chinese LungSCeQTL cohort and the European cohorts per cell type, with colored dots indicating eQTLs that replicate at a *P*-value less than 0.05. The direction of the Z score denotes the direction of the allelic effect with respect to the reference allele.

We further compared our sc-eQTL dataset with a recently published lung sc-eQTL study, which primarily focused on individuals with pulmonary fibrosis from European populations^21^. Overall, 35.5% of the sc-eQTLs identified in our dataset were reproducibly validated in corresponding cell types in the European cohort, with reproducibility ranging from 19.4% in B cells to 61.0% in ciliated cells (**Figure 3E**. Notably, rs1805 exhibited a significant and specific correlation with *MPZL2* expression in MP cells across both datasets, whereas rs929058 showed a population-specific association with *ROS1* expression in alveolar epithelial cells (AT2/AT1 cells), reaching statistical significance only in the Chinese cohort. These discrepancies may be attributed to the high proportion of fibrotic lung tissue in the European cohort, as well as potential ethnic disparities between the studies^49^.

Finally, we integrated the eSNPs identified in our sc-eQTL analysis with chromatin accessibility peaks derived from snATAC-seq profiling of lung tissue^32^. On average, 57.1% of sc-eQTLs localized within ± 5KB of the identified chromatin accessibility peaks of corresponding cell types, with proportions ranging from 23.1% in MyoF cells to 75.1% in Club cells. For example, rs929058, which regulates *ROS1* expression, mapped to an open chromatin region in AT2 cells. Notably, eSNPs in sc-eQTLs were significantly enriched near open chromatin peaks across various cell types compared to randomly selected SNPs **(Extended Data Figure 1**), highlighting the regulatory relevance of these genetic variants.

### Using sc-eQTL to Identify Causal Genes from Lung Cancer Susceptibility Loci

To enhance our understanding of lung cancer GWAS loci, we utilized the lung sc-eQTL database to identify candidate causal genes. By integrating data from five recent lung cancer GWAS studies^5–9^, we selected 33 independent genetic variants across 31 loci (MAF > 0.01, *P* < 0.05 in our Chinese GWAS) for further analysis. After integrating our sc-eQTL, bulk eQTL, and lung snATAC-seq databases^32^ with the GWAS datasets^8^, we identified 113 candidate causal genes, categorized into four evidence levels (**Methods**): (1) Level 1: 28 genes at 16 loci with significant eQTLs (*P*_fdr_ < 0.05) and colocalization signals (posterior probability 4 [PP4] ≥ 0.7); (2) Level 2: 17 genes at 5 loci with significant eQTLs but no colocalization signals; (3) Level 3: 64 genes at 7 loci with nominal association of eQTLs (*P* < 0.05); and (4) Level 4: 3 genes at 3 loci with only chromatin accessibility peaks from snATAC-seq (**Figure 4A**).

**Figure 4.**
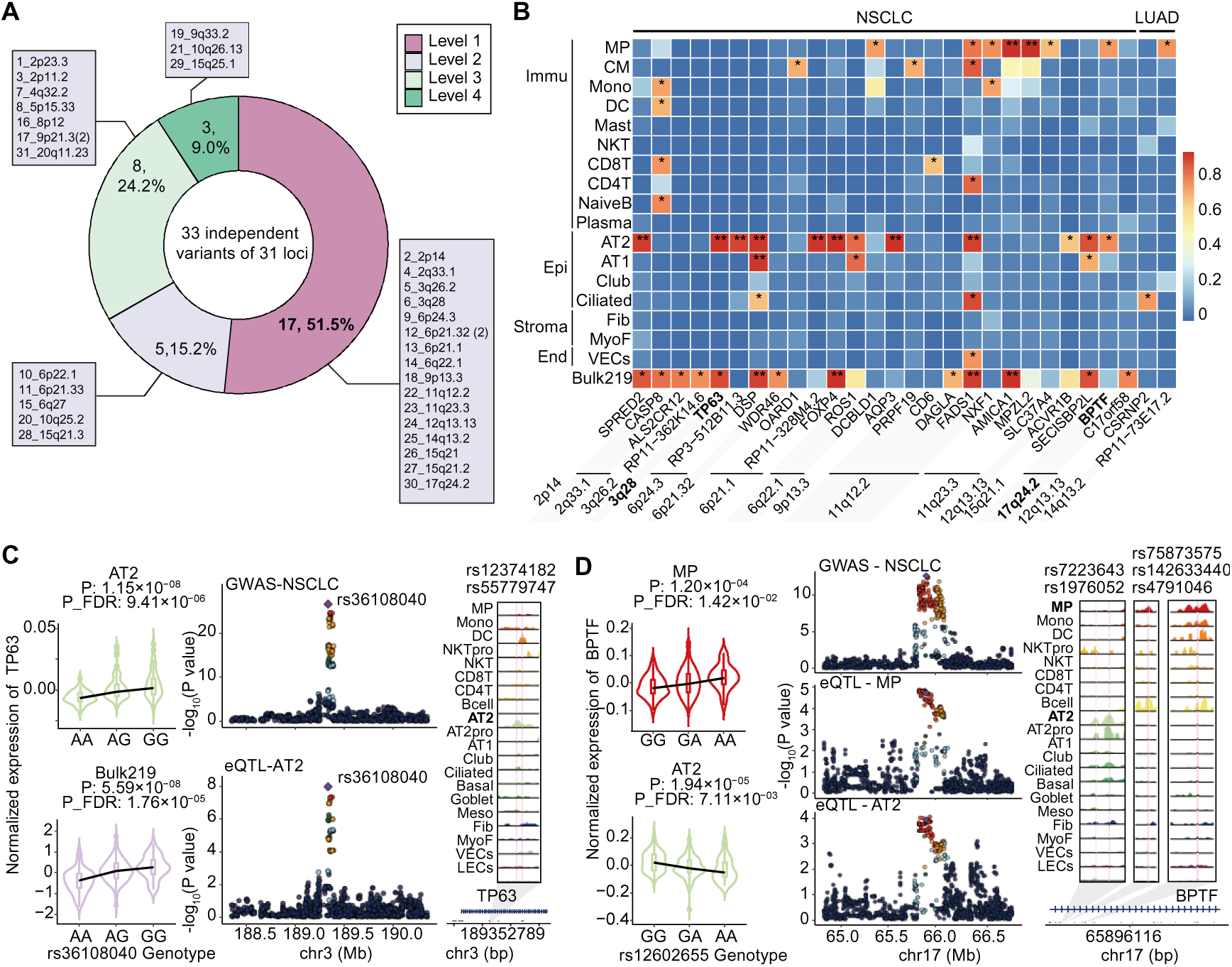
Dissection of identifying target susceptibility genes in the lung cancer GWAS loci. **(A)** Pie chart presents the details of annotating reported lung cancer GWAS loci by combining our eQTL database with lung snATAC-seq database. **(B)** Heatmap shows the PP4 of co-localization for target susceptibility genes in the lung cancer GWAS loci that colocalized with GWAS associations for NSCLC, LUAD, and LUAC. PP4 > 0.7 and PP4 > 0.95 are marked ‘*’ and ‘**’, respectively. **(C-D)** A detailed presentation examples of co-localization signals in the lung cancer GWAS loci including TP63 with rs36108040 (C) and BPTF with rs12602655 (D). Allelic violin plots show the effects of lead variants on gene expression in specific cell types. The y axis represents normalized expression of candidate genes, and the x axis represents genotypes of the lead variants. Regional plots show the overlap of GWAS and eQTL associations, with the lead variant indicated by a purple diamond and other points are colored according to their linkage disequilibrium (LD) with the lead variant. Sequencing tracks represent chromatin accessibility for the lead variants and their high LD variants (r² > 0.5), marked by rsIDs and vertical pink lines. Each track represents the aggregated snATAC signal of each cell type, normalized by the total number of reads in the regions. Arrow depicts the transcriptional direction of TP63(C) and BPTF (D).

Among the 28 candidate causal genes in Level 1, we observed complex, cell-type-specific regulatory patterns of lung cancer-associated variants. First, nearly half (46.43%, 13/28) of the identified genes showed colocalization in epithelial cells, predominantly AT2 cells (**Figure 4B**,). For example, the A allele of rs36108040 at 3q28 was significantly associated with decreased *TP63* expression in both AT2 cells (β = - 0.006, *P*_eQTL_=1.15×10^-8^, PP4=0.99) and bulk data (β =-0.407, *P*_eQTL_ = 5.59×10^-8^, PP4=0.94). Additionally, its high LD SNP rs55779747 (r^2^=0.86) was located within an AT2-specific snATAC peak (**Figure 4C**). Second, a subset (32.14%, 9/28) of genes exhibited colocalization signals exclusively in immune cells, suggesting that these variants may influence lung cancer susceptibility via the lung microenvironment. For instance, the T allele of rs6731171 was linked to an increased lung cancer risk and reduced *CASP8* expression in Mono, DC, CD8⁺ T, and Naïve B cells **(Extended Data Figure 2**). Third, we identified opposing colocalization signals for *BPTF* at 17q24.2 in MP and AT2 cells. The rs12602655-A allele was associated with increased *BPTF* expression in MP cells but decreased expression in AT2 cells. Further analysis revealed distinct genetic variants in high LD (r² > 0.5) with rs12602655 located within cell-type-specific snATAC peaks of MP and AT2 cells, respectively (**Figure 4D**). Finally, a broad regulatory pattern was observed for *FADS1* at 11q12.2 across multiple cell types. The rs174561-C allele was associated with decreased *FADS1* expression in MP, CM, AT2, AT1, and Ciliated cells, while increased its expression in CD4^+^ T cells **(Extended Data Figure 3**).

In addition to mapping candidate genes to specific cell types, we found that lung cancer-associated variants can simultaneously regulate multiple genes across different cell types. Nearly a quarter (25.81%, 8/31) of lung cancer susceptibility loci were colocalized with two or more candidate susceptibility genes across various cell types. A striking example is rs174561 at 11q12.2, which simultaneously regulates *FADS1* expression in six cell types, *NXF1* expression in MP and Mono cells, *PRPF19* expression in CM cells, CD6 expression in CD8^+^ T cells. Further analysis revealed that SNPs in high LD (r² > 0.5) with rs174561 were located at several distinct snATAC peaks, many of which were shared across different cell types **(Extended Data Figure 3**). Parallel patterns emerged at 11q23.3, where rs17121871 co-regulated *MPZL2* and *AMICA1* in MP cells, with SNPs in high LD (r² > 0.5) located at snATAC peaks specific to immune cells **(Extended Data Figure 4**). These multi-gene regulatory hubs, detectable only through cell-type-resolved analyses, underscore the necessity of sc-eQTL analysis in unraveling the genetic mechanisms underlying lung cancer susceptibility.

### Transcriptome-wide association analysis reveals novel lung cancer susceptibility genes

To identify additional lung cancer susceptibility genes at the cellular level, we further conducted cell type-specific transcriptome-wide association studies (TWASs). A total of 250 genes were identified with a significance of *P*_fdr_ < 0.1 across 17 lung cell types, including 178 genes in NSCLC, 163 genes in lung adenocarcinoma (LUAD) and 51 genes in lung squamous carcinoma (LUSC). Subsequent colocalization analysis of TWAS-identified candidates identified 30 lung cancer susceptibility genes across 25 loci, comprising 18 genes in NSCLC, 19 genes in LUAD, and 8 in LUSC (**Figure 5A**). Further integration with the lung snATAC-seq database^32^ revealed that 79.3% (23/29) of the eSNPs or their high LD variants associated with 24 genes were located within the snATAC-seq peaks of the corresponding cell types (**Figure 5A**).

**Figure 5.**
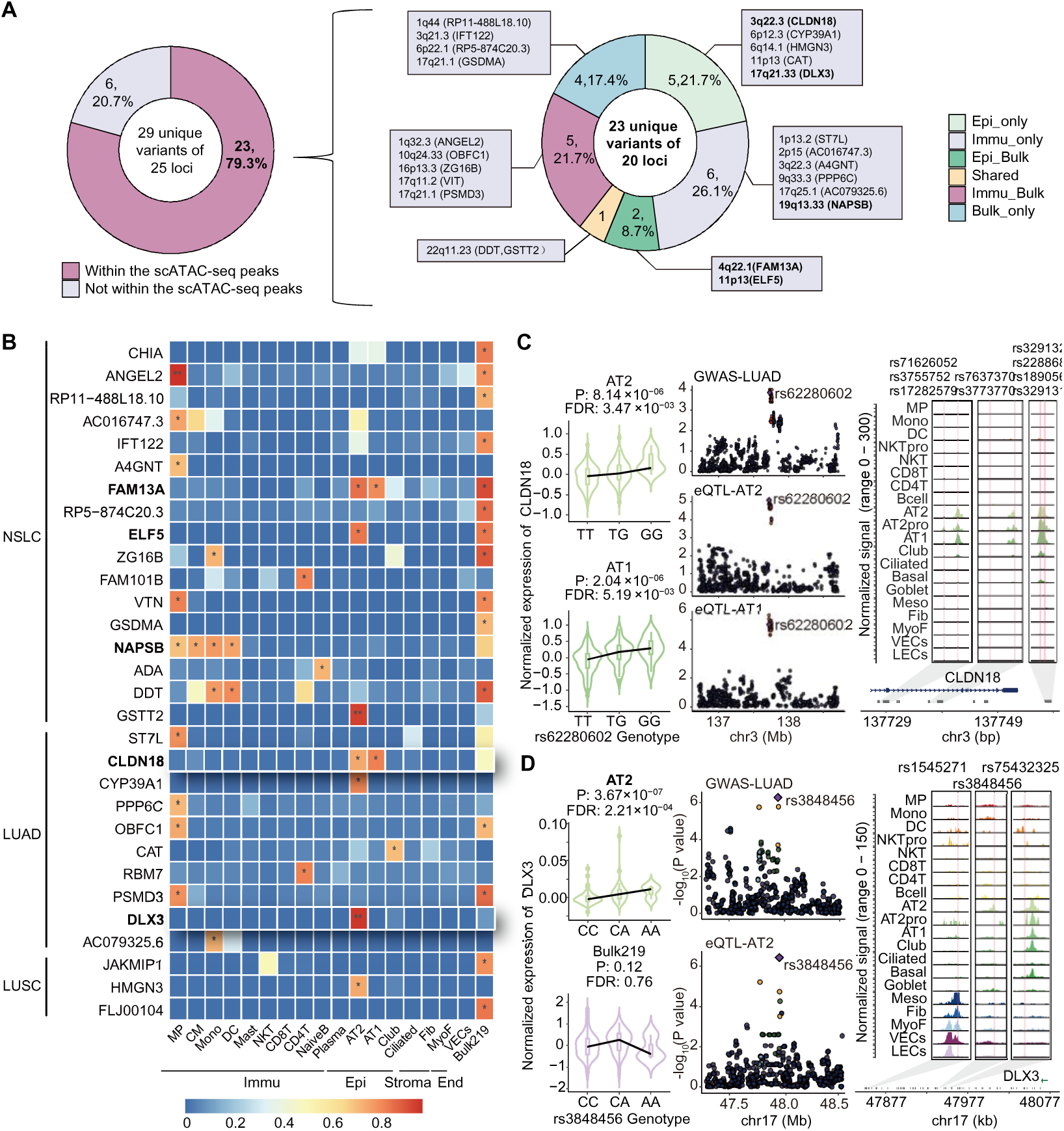
Dissection of identifying target susceptibility genes in novel lung cancer susceptibility oci. **(A)** Pie chart presents the details of the novel lung cancer susceptibility loci which were correlated with changes in gene expression levels and exhibited chromatin accessibility in corresponding cell types. These loci are further classified based on the functional mode of action of epithelial and immune populations. **(B)** Heatmap shows the PP4 of co-localization for target susceptibility genes in novel lung cancer GWAS loci that colocalized with GWAS associations for NSCLC, LUAD, and LUSC (*PP4 > 0.7, **PP4 > 0.95). **(C-D)** Detailed presentation of co-localization signals in novel lung cancer GWAS oci including CLDN18 with rs62280602 (C) and DLX3 with rs3848456 (D). Allelic violin plots show he effects of lead variants on gene expression in specific cell types. Regional plots show the overlap of GWAS and eQTL associations, with the lead variant indicated by a purple diamond. Other points are colored by LD with the lead variant. Sequencing tracks show chromatin accessibility for the lead variants and their high LD variants (r² > 0.5), marked by rsIDs and vertical pink lines. Each track represents the aggregated snATAC signal of each cell type, normalized by the total number of reads in he regions. Arrow depicts the transcriptional direction of CLDN18 (C) and DLX3 (D).

Approximately 50.0% (12/24) of the newly identified susceptibility genes were colocalized in immune cells, predominantly in MP cells (**Figure 5B**). Notably, we identified colocalization signals for *NAPSB* at 19q13.33 across multiple immune cell types, including MP, CM, Mono, and DC cells **(Extended Data Figure 5**). The lead SNP rs7247907 was consistently associated with increased lung cancer risk in both Chinese (OR=1.09, *P*=4.11×10⁻⁴) and European populations (OR=1.04, *P*=1.80×10⁻²)^5^. Importantly, prior evidence positions NAPSB as a molecular marker characterizing immunologically active tumor microenvironments^50^.

Additionally, eight novel lung cancer susceptibility genes were identified from epithelial cells, predominantly in AT2 cells, which was major cell type responsible for human lung adenocarcinoma^51,52^. Notably, *FAM13A* at 4q22.1 and *ELF5* at 11p13, previously implicated in chronic obstructive pulmonary disease^53^ and severe COVID-19^54^, respectively, were also found to be associated with lung cancer risk in TWAS with strong colocalization signals. Furthermore, rs62280602-T allele at 3q22.3 demonstrated significant associations with an increased risk of LUAD (OR=1.09, *P*=1.35×10⁻⁴) and decreased *CLDN18* expression in both AT1 (*P*_fdr_=5.19×10⁻³) and AT2 cells (*P*_fdr_=3.47×10⁻³). SNPs in high LD with rs62280602 localizes within chromatin accessibility regions specific to AT1 and AT2 cells (**Figure 5C**). *CLDN18* encodes a tight junction protein highly expressed in the lung, and Cldn18⁻/⁻ mice have been observed to exhibit abnormally large lungs due to increased AT2 cell proliferation and a higher frequency of lung adenocarcinomas^55^. Further validation in European populations^5^ confirmed consistent associations for 6/7 loci (86%), underscoring their relevance across populations.

Among the loci validated in both populations, *DLX3* at 17q21.33, a novel gene implicated in lung cancer, showed strong colocalization signals exclusively in AT2 cells (*P*_TWAS_=6.85×10⁻⁵, PP4=0.97). The rs3848456-A variant was significantly associated with an increased risk of LUAD in both the Chinese population (OR=1.17, *P*=5.12×10⁻⁷) and European population (OR=1.12, *P*=1.16×10⁻²), coupled with elevated *DLX3* expression in AT2 cells (**Figure 5D**). Further functional analysis indicated that rs3848456 variant was located within AT2-specific chromatin accessibility regions (**Figure 5D**), and luciferase reporter assays indicated that rs3848456-A conferred higher enhancer activity compared to rs3848456-C in AT2 cells (**Figure 6A**). Then, the functional role of *DLX3* was investigated by perturbing its expression through siRNA-mediated knockdown and plasmid-mediated overexpression in AT2 cells (**Figure 6B**). MTT assays revealed that downregulation of *DLX3* significantly inhibited cell viability, while overexpression of *DLX3* enhanced it (**Figure 6C**). Additionally, *DLX3* knockdown reduced colony formation capacity, while its overexpression promoted it (**Figure 6D**). Specially, knockdown of *DLX3* in AT2 cells increased the expression of AT2-to-AT1 transition markers (*SFTPA1*, *SFTPC*, *LAMP3*) and decreased the expression of LUAD plasticity markers (*MDK*, *HSF1*, *CEBPG*). In contrast, *DLX3* overexpression in AT2 cells upregulated LUAD plasticity markers, suggesting a phenotypic shift of AT2 cells towards a malignant state (**Figure 6E**).

**Figure 6.**
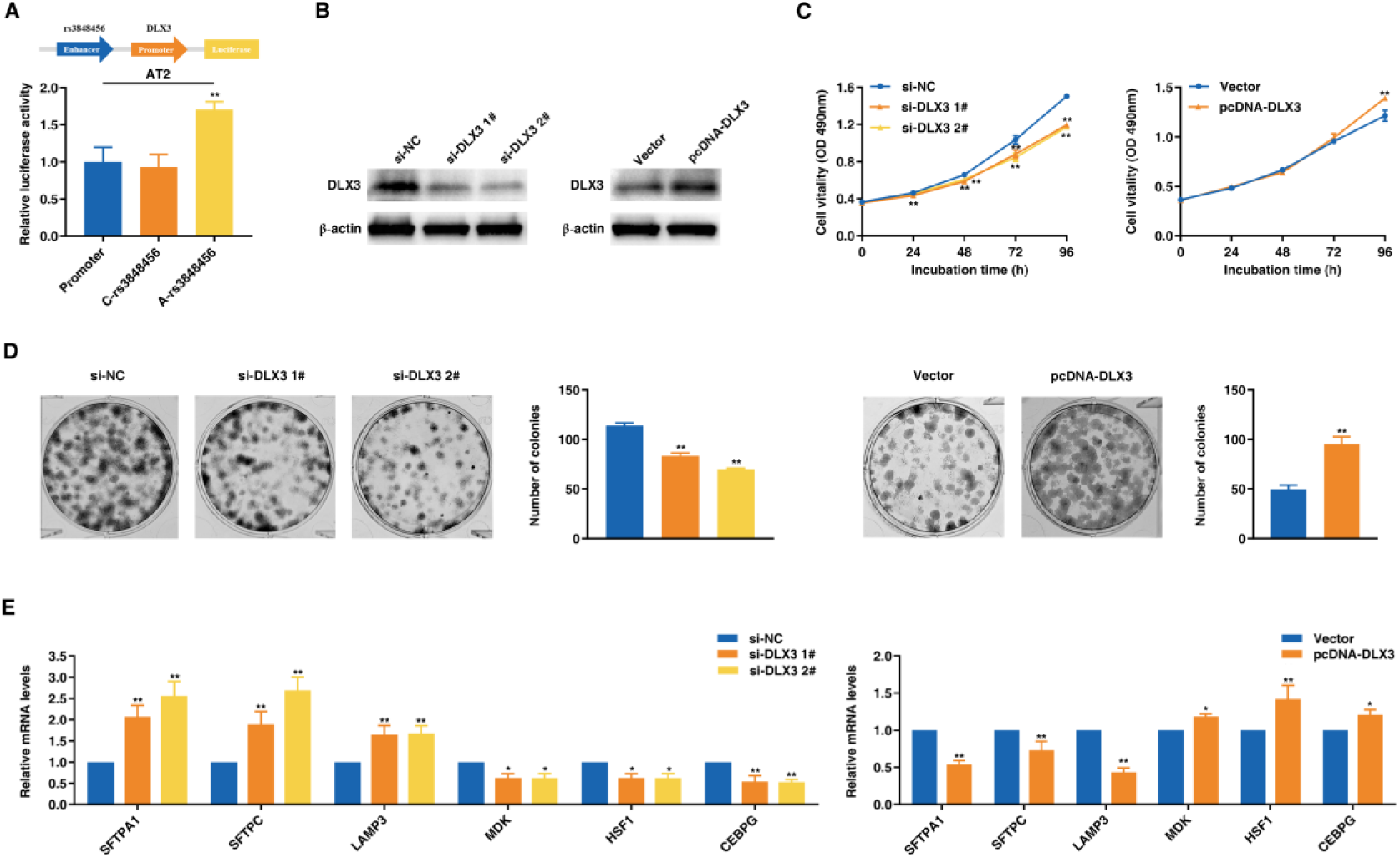
Functional mechanisms of DLX3 in AT2 cells. (A) Effect of DLX3 on the luciferase activity of the constructs containing the rs3848456[C] and the rs3848456[A] in AT2 cell. (B) Western blot assays detected the protein level of after knockdown and overexpression of DLX3. (C) MTT assays were performed to detect the cell viability of AT2 cell after DLX3 knockdown and overexpression. (D) Colony formation assays were performed to detect cell proliferation of AT2 cell after DLX3 knockdown and overexpression. (E) QRT-PCR assays detected the expression of related genes after knockdown and overexpression of DLX3. *P < 0.05 and **P < 0.01.

Interestingly, we also found that rs4822455 at 17q21.1 was colocalized with the *DDT* expression in Mono and DC cells, and *GSTT2* expression in AT2 cells simultaneously **(Extended Data Figure 6**). The lead SNP rs4822455 was nominally associated with lung cancer risk in Chinese GWAS (OR = 1.10, *P* = 8.15 × 10⁻⁸), and its high LD SNPs localizes within several distinct chromatin accessibility regions. *DDT* encodes a cytokine similar to macrophage migration inhibitory factor, which binds to CD74 with high affinity, activating the ERK1/2 MAP kinase and proinflammatory pathways^56^. Meanwhile, *GSTT2* encodes a detoxifying enzyme responsible for inactivating reactive oxygen species and reducing DNA damage^57^. These findings highlight the pleiotropic effects of genetic variants in distinct cell types, collectively contributing to lung cancer susceptibility with the intricate regulatory patterns.

### Druggable targets identified from colocalization results

To bridge genetic architecture and clinical therapeutic translation, we systematically evaluated the druggability of NSCLC candidate genes identified in this study using Drug-Gene Interaction Database^58^. Among these, 32.1% (9/28) of genes in established susceptibility regions and 20.8% (5/24) in novel loci had interactions with either approved drugs or those under development, suggesting their potential as clinically actionable targets. For instance, *ROS1*, a receptor tyrosine kinase frequently mutated in cancers, is a key target for over 20 approved drugs, including crizotinib and entrectinib, widely used in NSCLC treatment. Strikingly, this study highlights two novel genetic signatures as promising drug targets: (1) Zolbetuximab, a first-in-class chimeric immunoglobulin G1 monoclonal antibody highly specific for CLDN18.2, is in phase III trials (NCT01630083) for treating CLDN18.2-positive gastric cancer and presents potential for lung cancer applications; (2) IXAZOMIB, a clinically approved receptor inhibitor of DLX3, is currently used for treating a variety of malignant tumors and may be repurposed for lung cancer therapy. These findings underscore the therapeutic potential of candidate genes identified in this study, paving the way for drug development and repurposing.

## Discussion

In this study, we present a comprehensive profiling of regulatory genetic variants across major lung cell types in the Chinese population, leveraging scRNA-seq combined with a pooled multiplexing strategy to identify cell type-specific eQTLs. We identified 16,785 independent eQTLs across 17 cell types, mapping to 8,708 eGenes, with full summary statistics accessible via our online database (http://ccra.njmu.edu.cn/LungSCeQTL). Notably, the majority of sc-eQTLs and over half of the eGenes exhibited cell type-specific regulatory effects, underscoring the critical role of cellular context in genetic regulation. Integrating sc-eQTLs with lung cancer risk loci identified through GWAS, we pinpointed putative causal genes and the specific cell types through which they exert their pathogenic effects. Additionally, TWAS and colocalization analyses uncovered novel susceptibility genes, expanding the spectrum of genetic factors implicated in lung cancer. Together, these findings provide new insights into cell type-specific genetic regulation in the lung, advancing our understanding of lung cancer susceptibility and laying the groundwork for precision medicine strategies targeting cell-specific genetic pathways.

By integrating macroscopic environmental and behavioral factors with population-scale single-cell transcriptomic profiles, our analysis revealed significant impacts of various macro-factors on lung cellular composition. Among them, smoking and chronic respiratory diseases (CRD) emerged as the most influential in shaping lung tissue cellular architecture. Specifically, smoking was associated with the expansion of epithelial cells, such as AT2 and AT1, likely due to their role in lung repair following smoking-induced injury^59,60^. Additionally, consistent with previous studies showing that airway inflammation-driven CRD, such as COPD, are characterized by vascularization, lymphangiogenesis, and the expansion of supporting cells^61^, our findings demonstrated a higher proportion of epithelial cell, myofibroblasts and lymphatic endothelial cells in patients with CRD.

Compared to conventional bulk-eQTL analyses, sc-eQTL analysis offers several advantages in mapping allele-specific transcriptional regulation. Consistent with prior findings^21–25^, we observed that a substantial proportion of sc-eQTLs (77.2%) were undetected in matched bulk-eQTL datasets. This discrepancy was particularly pronounced in low-abundance cell types, highlighting that single-cell resolution enables the identification of unique cell-specific regulatory mechanisms that are often obscured in bulk approaches, especially in the context of rare cell-type-specific genetic regulation. Moreover, in line with previous studies^21,62^, sc-eQTLs were found to be located farther from transcription start sites (TSS) and enriched in distal regulatory elements, such as enhancers—particularly for cell type-specific sc-eQTLs. This suggests that sc-eQTL analysis facilitates the detection of enhancer-mediated regulation that may be overlooked in bulk analyses.

Furthermore, compared to existing lung sc-eQTL databases derived from European populations^21^, we constructed an sc-eQTL database using a larger cohort of normal lung tissues. Despite applying more stringent statistical thresholds, we identified 21.7% more eGenes. Notably, while Natri et al.^21^ reported that only 6.5% of sc-eQTLs exhibited cell type specificity, our study revealed that over 76% were cell type-specific—a finding consistent with single-cell eQTL studies in PBMCs, brain tissue, and colorectal tissues^22,25,63^. This discrepancy may be attributed to the dysregulated cellular circuitry and altered cell proportions in whole lung tissue caused by pulmonary fibrosis^35,40^. Thus, our study not only provides a more comprehensive sc-eQTL database for a larger normal lung sample size but also serves as a valuable resource for investigating single-cell-level genetic regulation across diverse genetic backgrounds, extending beyond pulmonary fibrosis to other chronic lung diseases.

This study further explored the association between sc-eQTLs and lung cancer genetic susceptibility. Through colocalization analysis, we identified 28 colocalized genes from 17 known NSCLC susceptibility regions in GWAS and determined their potential cellular targets. Notably, nearly half (13/28) of these causal genes were detectable only at the single-cell level, underscoring the advantage of sc-eQTL analysis in identifying potential NSCLC causal genes. Furthermore, we observed the most significant enrichment of GWAS risk loci in AT2-eQTL and MP-eQTL, with 12 and 9 colocalized genes specifically identified in these cell types, respectively. This finding suggests that AT2 and MP cells may serve as key cellular mediators of NSCLC susceptibility, aligning with previous reports that macrophage-driven dedifferentiation of AT2 cells into progenitor-like states may synergistically promote carcinogenesis^35,64^.

By integrating TWAS and COLOC analyses to refine the identification of risk genes, we uncovered 24 novel candidate causal genes. Several of these loci, including *FAM13A*, *ELF5*, *CLDN18*, and *CAT*, have been previously implicated in pulmonary disorders^53–55,65–67^. Furthermore, our systematic analysis and *in vitro* experiments confirmed *DLX3* at 17q21.33—a homeobox transcription factor involved in differentiation and cell cycle regulation^68–70^—as a novel factor influencing lung adenocarcinoma through its specific regulatory effects in AT2 cells. Notably, more than 20% of these newly identified NSCLC susceptibility genes represent potential druggable targets, with over 37% showing significant associations with lung cancer in European populations^5^. These findings underscore the translational potential of NSCLC susceptibility gene discoveries based on our sc-eQTL database.

Notably, we identified a prevalent pattern of cell-type-specific pleiotropic genetic regulation (47%, 8/17) in established NSCLC susceptibility regions. One scenario involves an LD block within a single lung cancer susceptibility locus regulating multiple genes within the same cell type. For instance, the 11q23.3 locus simultaneously modulates *AMICA1*, *MPZL2*, and *SLC37A4* expression in MP cells. Additionally, we discovered that lung cancer susceptibility loci can regulate the same gene across different cell types, exhibiting bidirectional regulation^25^. A representative example is the pleiotropic eQTL rs12602655 at 17q24.2, which exhibits bidirectional regulation of BPTF in AT2 versus MP cells. *BPTF*, a bromodomain PHD finger transcription factor essential for chromatin remodeling and transcriptional regulation, has been reported to promote lung carcinogenesis through mechanisms including angiogenesis activation and T cell immunity suppression^71–75^. We also identified instances where a single susceptibility locus regulated distinct genes in different cell types. For example, *DCBLD1* and *ROS1* at 6q22.1—clinically validated therapeutic targets in lung cancer^76–79^—are regulated in MP and AT2 cells, respectively. This pleiotropic regulatory pattern at lung cancer susceptibility loci may arise from distinct subsets of high-LD trait-associated variants that localize to different regulatory elements across different cell types^32^. However, we found that half of these pleiotropic regions share regulatory elements across multiple cell types, suggesting that cell-type-specific transcription factor expression and chromatin spatial conformation may also contribute to this phenomenon^80–82^. Collectively, these insights further underscore the value of single-cell eQTL analysis in unraveling the complexity and cellular-context dependency of lung cancer susceptibility.

While our study provides key insights into the genetic regulation of lung cell types, several limitations should be acknowledged. First, although this represents the largest single-cell eQTL study of lung tissue to date, the sc-eQTL analysis was constrained by the limited number of cells captured for rare cell types. Future efforts should prioritize enriching rare cell types to comprehensively delineate their regulatory landscape. Second, the functional validation of *DLX3* and its potential role in lung cancer development was conducted exclusively through *in vitro* experiments. The roles of candidate genes in lung cancer pathogenesis require further confirmation in in vivo models and clinical cohorts. Third, our cohort primarily consisted of individuals of Chinese ancestry. While we integrated single-cell omics data from a Korean population for functional validation and replicated some findings in European populations, additional multi-ethnic studies are necessary to ensure the broader generalizability of our results.

In summary, this study provides the first comprehensive atlas of lung cancer susceptibility genes at single-cell resolution, revealing cell type-specific regulatory mechanisms and novel susceptibility genes. By integrating multi-omics data with functional validation, we advance the understanding of the genetic and cellular basis of lung cancer. These findings not only offer critical insights for elucidating the impact of heterogeneous genetic backgrounds on pulmonary pathologies but also pave the way for precision medicine approaches targeting specific cell types and genetic pathways.

## Methods

### Ethics statement

The present study was approved by the Institutional Review Board of Nanjing Medical University, and written informed consent was obtained from all participants prior to study enrollment.

### Participants, samples collection and tissue processing

The Chinese LSCQ cohort comprised 226 participants who underwent thoracic surgery at Nanjing Chest Hospital between October 2021 and July 2022. Each participant completed a structured baseline interview.

To minimize recruitment bias, lung parenchyma tissue samples were collected from fresh specimens following surgical resection of lesions, including (i) adjacent non-malignant lung tissue (>5 cm from the tumor edge) and (ii) normal lung tissue from non-neoplastic lung diseases. For single-cell RNA sequencing (scRNA-seq), fresh lung tissues were rinsed with PBS and immersed in sCelLive Tissue Preservation Solution (Singleron) on ice. After three washes with HBSS, the tissue was minced and digested in 3 mL sCelLive Tissue Dissociation Solution (Singleron) at 37°C for 15 minutes using the Singleron PythoN Tissue Dissociation System. The resulting cell suspension was filtered through a 40-μm sterile strainer, and red blood cells were lysed using GEXSCOPE red blood cell lysis buffer (Singleron) at room temperature for 5-8 minutes. After centrifugation (300 × g, 5 minutes, 4°C), the cell pellet was resuspended in PBS, and cell concentration and viability were assessed using a TC20 automated cell counter (Bio-Rad). Samples were processed into 29 pools including six replicates, with each pool containing cells from eight donors. A separate lung tissue sample was stored in 1 mL RNAlater and snap-frozen in liquid nitrogen. Peripheral blood samples were collected pre-surgery, processed within 4 hours, and centrifuged at 3,000 × g for 5 minutes at room temperature. Plasma, leukocytes, and red blood cells were aliquoted into cryotubes and stored at −80°C and −20°C, respectively.

### Genotyping, imputation and quality control

Genotyping was performed using the Infinium Asian Screening Array (ASA) with ∼670,000 markers following the manufacturer’s protocol. Quality control (QC) included filtering for SNP and individual call rates >95%, Hardy-Weinberg equilibrium (HWE) p-value >0.001, and minor allele frequency (MAF) >0.001. After QC, 530,011 SNPs from 226 individuals were retained. Genotype data were phased using SHAPEIT v2.12^80,81^, and imputation was performed using IMPUTE2 v2.3.1^82^ with the 1000 Genomes Project Phase III reference panel. We retained 4,838,930 variants with imputation quality INFO > 0.8 and MAF > 0.01 for cis-eQTL analyses.

### Single-Cell RNA Sequencing and Raw Data Preprocessing

Single-cell suspensions at concentrations range of 2.48 × 10⁵-4.50 × 10⁵ cells/mL in PBS were loaded onto a microwell chip using the Singleron Matrix Single Cell Processing System. Barcoding beads were retrieved from the microwell array, and captured mRNA was reverse-transcribed into cDNA, amplified via PCR, fragmented, and ligated with sequencing adapters. scRNA-seq libraries were prepared with the GEXSCOPE Single Cell RNA Library Kit (Singleron) protocol^83^. Libraries were diluted to 4 nM, pooled and sequenced on an Illumina NovaSeq 6000 to generate 150 bp paired-end reads.

Raw gene expression matrices were processed using Singleron’s Celescope software v1.1.8. QC and adapter trimming were performed with FASTQC v0.11.7 and Cutadapt v1.17, including poly-A tail and adapter sequence removal. Cell barcodes and unique molecular identifiers (UMIs) were extracted, and reads were aligned to the GRCh37/hg19 genome using STAR v2.6.1a^84^. UMI and gene counts were quantified with featureCounts v2.0.1^85^ to construct the expression matrix.

### Demultiplexing scRNA-seq pools

Donor assignment was performed using Demuxlet^33^, which employs a maximum likelihood approach to match cell-droplet genotypes (from 3’ RNA-seq reads) to imputed donor genotypes. Cell-droplets were classified as singlets (matching a single donor), doublets (matching multiple donors), or ambiguous (insufficient genotype information). Singlet cell-droplets were assigned to donors based on genotype posterior probabilities using 2,592,947 exonic variants. Only successfully assigned singlets were retained for downstream analysis.

### Quality control, cell-type clustering, and cell-type identification

Downstream analysis of scRNA-seq data was performed using the Seurat R package v4.2.0^86^. Cells were filtered based on the following criteria: UMIs counts between 500 and 50,000, gene counts between 200 and 6,000, and mitochondrial content >50%. After filtering, 845,976 cells were retained. Cells were then assigned to donors using the Demuxlet, and non-singlet cells, cells with low counts in duplicates, and doubles were removed. For individuals, pairwise relatedness was estimated using PLINK v1.9^87^ and individuals with poor genotype data or low cell counts were excluded. Finally, we retained 562,255 cells from 222 individuals for further analysis. Data were normalized and scaled using the NormalizeData and ScaleData. The top 2000 variable genes were selected with the FindVariableFeatures, followed by principal component analysis (PCA) using RunPCA and clustering using the top 50 principal components (FindClusters, resolution = 0.3). Differentially expressed genes (DEGs) between clusters were identified using FindAllMarkers. UMAP was used for 2D visualization of cell clusters.

The resulting clusters were divided into 7 major cell subgroups based on marker gene expression: Myeloid cells, mast cells; T cells; B cells; epithelial cells; stroma, and endothelial cells (ECs). Cells expressing markers for multiple cell types were classified as double-marker cells and excluded. Further classification of cell types was refined using both unsupervised and supervised clustering.

### Testing cell types for association with macroscopic factors

We used MASC v0.1^41^, a single-cell cluster association mixed effect model, to assess associations between cell types and potential lung cancer risk factors^43^, accounting for confounders at both the cell and donor levels. For each factor, we performed likelihood ratio tests (LRT) and used a gamma-distributed test statistic to evaluate the P-value distribution, considering factors significant if gamma P < 0.05. Additionally, we used linear regression with the “Relaimpo” R package (v2.2.6) to determine the contribution of significant factors to the variation in cell type proportions, based on their relative importance.

### Single-cell cis-eQTL mapping and conditional eQTL analysis

eQTL mapping was performed using fastQTL^46^ following the GTEx pipeline^18^. Linear regression was used to test for associations between SNP genotypes within a 1-Mb cis-region (including the gene body) and gene expression levels for each cell type, adjusting for covariates. For each cell type, we calculated the mean expression of each gene per donor using normalized counts, excluding genes expressed in <1% of cells and donors with <5 cells from that cell type. SNP with call rates <95%, HWE *p*-values <0.001, and MAF <0.05 were filtered out. Ultimately, cis-eQTL mapping was performed for 17 cell types.

For each cell type, we included eight macroscopic factors influencing lung tissue composition as covariates. To correct the potential influence of population structure and unmeasured factors in gene expression, we calculated the first 10 genotype principal components (PCs) and the first 15 PEER. We sequentially tested the effects of 1-10 PCs and 1-15 PEER factors as covariates in the linear regression model for cis-eQTL analysis of AT2 cells and observed an increase in the number of eGenes after adding the first PEER factor. Therefore, for each cell type, we included sex, age, smoking metrics, BMI, area, tea drinking, education level, CRD history, the first 10 PCs, and the first PEER factor as covariates. eQTL significance was determined with a false discovery rate (FDR) threshold of 0.05.

To identify independent genetic variants, we performed conditional eQTL analysis^22^ for each cell types. The top SNP (eSNP1) for each eQTL was identified based on the smallest nominal *p*-value. A second round of analysis was conducted to identify the second most significant SNP (eSNP2) by regressing out eSNP1. No other independent signals were found after the second round, so the analysis was repeated up to two rounds.

### Sharing of cis-eQTL signal between cell types

To quantify the extent of cis-eQTL sharing across cell types, we conducted a joint analysis of FastQTL-derived effect sizes and their standard errors using multivariate adaptive shrinkage (MASHR v0.2.73) (https://stephenslab.github.io/mashr/articles/eQTL_outline.html)^48^. SNP-gene pairs were included in the sharing analysis if they were significant (local false sign rate [lfsr] < 0.05) in at least one of the cell types. eQTL effects were considered shared in sign if they exhibited the same directionality across cell types and shared in magnitude if their effect sizes differed by no more than a factor of two.To determine whether each eGene was associated with distinct variants across cell types, we examined 272 pairs of cell types with shared eGenes. For cases where a gene was linked to different eSNPs in multiple cell types, we conducted a conditional analysis to assess the impact of the primary eSNP’s effect size after adjusting for the secondary eSNP. (i) If the eSNPs tagged independent variants, the allelic effect of the primary eSNP should remain stable. (ii) If they tagged the same causal variant, the effect size of the primary eSNP should decrease. A significant change in P value (P ≥ 0.05 after conditional analysis) indicates a shared eQTL, whereas no significant change (P < 0.05) suggests that the eQTLs are distinct between cell types.

### Bulk RNA Extraction, Sequencing and quantification

Total mRNA was extracted from the adjacent lung tissues corresponding to the samples used for single-cell RNA sequencing. RNA quantity and integrity were assessed with the RNA Nano 6000 Assay Kit on the Bioanalyzer 2100 system (Agilent Technologies). A total of 219 samples met quality standards with matching eligible scRNA-seq data (Bulk219). PolyA+ mRNA was isolated using the Illumina TruSeq RNA Sample Preparation Kit and sequenced on the Illumina NovaSeq 6000 platform, generating paired-end 150 bp reads. Raw FASTQ files underwent quality control with FastQC v0.11.7, which included adapter trimming, removal of reads containing N bases, and filtering of low-quality reads. Reads were aligned to the GRCh37/hg19 genome using STAR v2.4.2a, and gene expression was quantified with FeatureCounts, based on GENCODE v19 (July 2013 freeze) annotations.

### Bulk RNA-seq cis-eQTL mapping

Cis-eQTL mapping was performed on Bulk219 using fastQTL^46^ following the GTEx pipeline^18^. SNPs within a 1 Mb cis-region around the transcription start site (TSS) were tested for associations with gene expression. Covariates included the first 10 PCs, sex, age, smoking metrics, BMI, area, history of tea drinking, education level, history of CRD, and the first 30 PEER factors. Significant SNP-gene pairs were identified using an FDR threshold of <0.05.

### Replication of cell-type cis-eQTLs in Bulk219 and previous studies

We assessed the replication of sc-eQTLs in the Bulk219 dataset by comparing the direction and significance of allelic effects. Conversely, we evaluated the replication of Bulk219 cis-eQTLs in sc-eQTLs to determine the number of cell types with overlapping eGenes and eSNPs. Additionally, for each cell type, we replicated our Chinese sc-eQTLs in the European lung sc-eQTL dataset^21^ by comparing directions and significance of allelic effect using Z-scores.

### Enrichment of cis-eQTLs in regulatory regions

To assess the enrichment of cis-eQTLs in regulatory regions, we utilized a recently published single-nucleus ATAC-seq (snATAC-seq) dataset from 15 donor lungs (GSE241468)^32^ Data were processed using Signac v1.9.0^88^ and Seurat v4.2.0^86^ with the GRCh37/hg19 reference. After QC, RNA counts were normalized using NormalizeData, and ATAC peaks were normalized with RunTFIDF. Latent Semantic Indexing (LSI) was applied to ATAC data for dimensionality reduction, and PCA was used for RNA data. Cell type identities were assigned based on canonical markersand cell type-specific ATAC peaks were identified using MACS2^89^. For each cell type, we calculated the genomic distance between each cis-eQTL and its nearest peak. Similarly, we computed distances between each tested SNP and its nearest peak. To assess significance, null distributions were generated with tenfold bootstrap replicates, and distances were compared using a t-test. Additional snATAC-seq data from two donor lungs (GSE214085)^90^ was also included in the analysis, following the same procedures. We also used AT2 cells ChIP-seq data from two donors (GSE150527)^91^ to map and annotate cis-eQTLs in regulatory element regions.

### Colocalization analysis

We performed Bayesian colocalization using the COLOC R package (v5.2.2)^92^ to integrate cis-eQTLs with lung cancer GWAS summary data. We conducted overall NSCLC and stratified analyses for LUAD and LUSC. The COLOC method tests whether a shared genetic variant affects both gene expression and disease phenotype by fitting eQTL and GWAS data into a model and calculating posterior probabilities. For each gene, pairwise tests were done within a 1 Mb window. Genes with a posterior probability of colocalization (PP4) > 0.7 were considered co-localized with GWAS results.

### Linking the lung cancer GWAS loci to target genes in 4 levels

We integrated data from five recent lung cancer GWAS studies (McKay_2017^5^, Dai_2019^6^, Byun_2022^7^, Wang_2022^8^, and Shi_2023^9^) across European, East Asian, and African ancestries, identifying 33 independent genetic variants across 31 loci (MAF > 0.01 in East Asian populations, p < 0.05 for lung cancer GWAS in China, and R² < 0.5 within ±250kb of other variants). We then combined our sc-eQTL, Bulk219 eQTL, and lung snATAC-seq databases^32^ with our Chinese NSCLC GWAS data^8^ to assign target genes for NSCLC GWAS loci in four levels:

- Level 1: Genes with significant eQTLs (FDR < 0.05), colocalization signals (PP4 ≥ 0.7), and chromatin accessibility peaks from snATAC-seq data
- Level 2: Genes with significant eQTLs (FDR < 0.05) but no colocalization signals
- Level 3: Genes with nominal eQTL associations (P < 0.05)
- Level 4: Genes with colocalizing chromatin accessibility peaks from snATAC-seq data

### Transcriptome Wide Association Study (TWAS)

TWAS was performed for each lung cell type using FUSION software^93^. We estimated gene expression heritability in each cell type using GCTA-GREML^94^. Gene with significant heritability (cis-hg2, p < 0.1) were considered for further analysis. Cell type-specific expression profiles were used as reference panels to compute gene expression weights via prediction models in FUSION (top1, lasso, blup, and enet). Expression weights were derived from SNPs within ±1 Mb of each gene and integrated with GWAS results to estimate the association between gene expression and lung cancer. Stratified analyses for LUAD and LUSC were also performed. Genes were deemed significant if they met an FDR < 0.1 and MODELCV P < 0.05. A similar analysis was conducted on the Bulk219 dataset.

### Validation of the novel NSCLC susceptibility loci in European populations

To verify the generalizability of the novel susceptibility loci identified in this study, we used the European populationsdatabase including 14,803 cases and 12,262 controls^5^ for validation.

### Cell culture

Primary human AT2 cells was purchased from ZQXZBIO (Shanghai, China). AT2 cells were cultured with DMEM/F12 medium (CORNING, NY, USA) supplemented with 10% FBS and 1% penicillin/streptomycin at 37 °C and 5% CO_2_.

### Cell transfection

According to the manufacturer’s instructions, the small interfering RNA (siRNA) was transfected into cells using Lipofectamine 2000 (Invitrogen, Thermo Fisher Scientific, Sunnyvale, CA, USA), including the corresponding controls. The DLX3 sequence was subcloned into a pcDNA3.1 vector (Invitrogen, Thermo Fisher Scientific, Sunnyvale, CA, USA). The pcDNA-DLX3 vector was transfected into cells using the X-tremeGENE HP DNA transfection reagent (Roche, Basel, Switzerland). The experiments were independently replicated three times.

### Western Blot assays

Transfected cells were lysed with RIPA (KeyGEN Biotech, Shanghai, China) mixed with protease inhibitors (MCE, NJ, USA) and centrifuged to extract proteins. Proteins were separated by electrophoresis in 4%-20% SurePAGE. After transferring the proteins on the gels to 0.22μm PVDF membranes (Millipore, Massachusetts, USA), then the membranes were incubated with specific antibodies overnight at 4°C. DLX3 antibody (Dilution ratio, 1:1000, A20948) were from Abclonal. After incubating the corresponding secondary antibody, the final protein bands were observed on a molecular imager (Bio-Rad, Hercules, USA).

### Dual-luciferase reporter assays

Enhancer activity was detected using a dual-luciferase reporter gene kit (Easybio, Bejing, China). The promoter plasmid was obtained by inserting the promoter regions of the DLX3 gene into the PGL3 vector, and the promoter-enhancer plasmid was obtained by inserting the mutated enhancer fragment into the promoter plasmid. Co-transfected the reporter gene plasmid and Renilla plasmid into the cells. For the transfection of AT2 cells, used fugene (Roche, Basel, Awit). After 48 hours, the cells were lysed with a kit and luciferase activity was measured. The experiments were independently replicated three times.

### Cell proliferation analysis

The detection of cell viability used the MTT assay kit (Biofroxx, Einhausen, GER). The treated cells were uniformly seeded in a 96-well culture plate at a certain concentration, and MTT reagent was added every 24 hours. After incubation in an incubator for 4 hours, dimethyl sulfoxide was added and the absorbance value at 490 nm was detected. In the colony formation experiment, the treated cells were inoculated in a 24-well culture plate. After the cells growed in the incubator for about 2 weeks, visible colonies can be formed. After fixation with methanol and staining with crystal violet, counting can be performed. The experiments were independently replicated three times.

### RNA extraction and qRT-PCR analyses

According to the manufacturer’s instructions, the total RNA extracted by TRIzol (Invitrogen, USA) was reverse transcribed into cDNA using a High Efficiency Reverse Transcription kit (Vazyme, Jiangsu, China). Then, qRT-PCR experiments were performed on the QuantStudio 7 Flex system using the TB GreenTM Premix Ex Taq^TM^ kit (Takara, Kusatsu, Japan). GAPDH was used as an internal reference in the experiment process.

### Druggable targets analysis

We investigated gene-drug interactions for 28 reported and 24 novel lung cancer susceptibility genes using the Drug-Gene Interaction Database (DGIdb v5.0, https://dgidb.org)^58^. We then searched the ChEMBL database (v34, https://www.ebi.ac.uk/chembl/)^95^ to identify drugs targeting these genes. Additionally, the DrugBank database (v5.1.12, https://go.drugbank.com/)^96^ was used to explore pharmacodynamics and mechanisms of action for potential drugs.

### Statistics and reproducibility

Statistical analyses were performed using R v.4.1.3, as detailed in the Methods and figure legends.

## Data availability

We developed a publicly accessible online platform at http://ccra.njmu.edu.cn/LungSCeQTL to facilitate exploration of eQTLs and eGenes across various lung cell types. Data sharing complies with the Ministry of Science and Technology of the People’s Republic of China and has been archived in the China Human Genetic Resources Bank (no. BF2025011116170).

## Code availability

The code used for the analysis presented in this study is available at https://github.com/LungHealthFu/LungSCeQTL_Analysis

## Acknowledgments

This work was supported by the Ministry of Science and Technology of China no.2024ZD0520000 (2024ZD0520003) to H.M.; the National Natural Science of China no.81922061 to H.M., grant no.82388102 to H.S., grant no. 82473708 to M.Z.; and the Excellent Youth Foundation of Jiangsu Province no. BK20220100 to M.Z. We are grateful to the participants and researchers who contributed to this study.

## Author contributions

Meng Zhu, Hongbing Shen and Hongxia Ma initiated, conceived, and supervised the study. Yating Fu and Meng Zhu developed the methodology, performed bioinformatics/statistical analyses and prepared the manuscript. Chen Jin, Linnan Gong, Yuanlin Mou Caochen Zhang and Shihao Wu assisted with data analysis, along with Yi Wang, Yating Fu, Chenying Jin, Chen Ji, Xinyuan Ge, Yahui Dai were involved in sample and clinical information collection. Chang Zhang, Jiaying Cai, Sunan Miao, Huimin Ma, Xiaoyang Ma, Mengping Wang and Erbao Zhang conducted the related experiments. Yating Fu and Lijun Bian created the website (LungSCeQTL), and Lijun Bian, Juncheng Dai, Zhibin Hu and Guangfu Jin contributed to data interpretation. All authors critically reviewed the manuscript. Meng Zhu, Hongbing Shen and Hongxia Ma acquired the funding.

## Competing interests

The authors declare no competing interests.

---

**Extended Data Figure 1.**
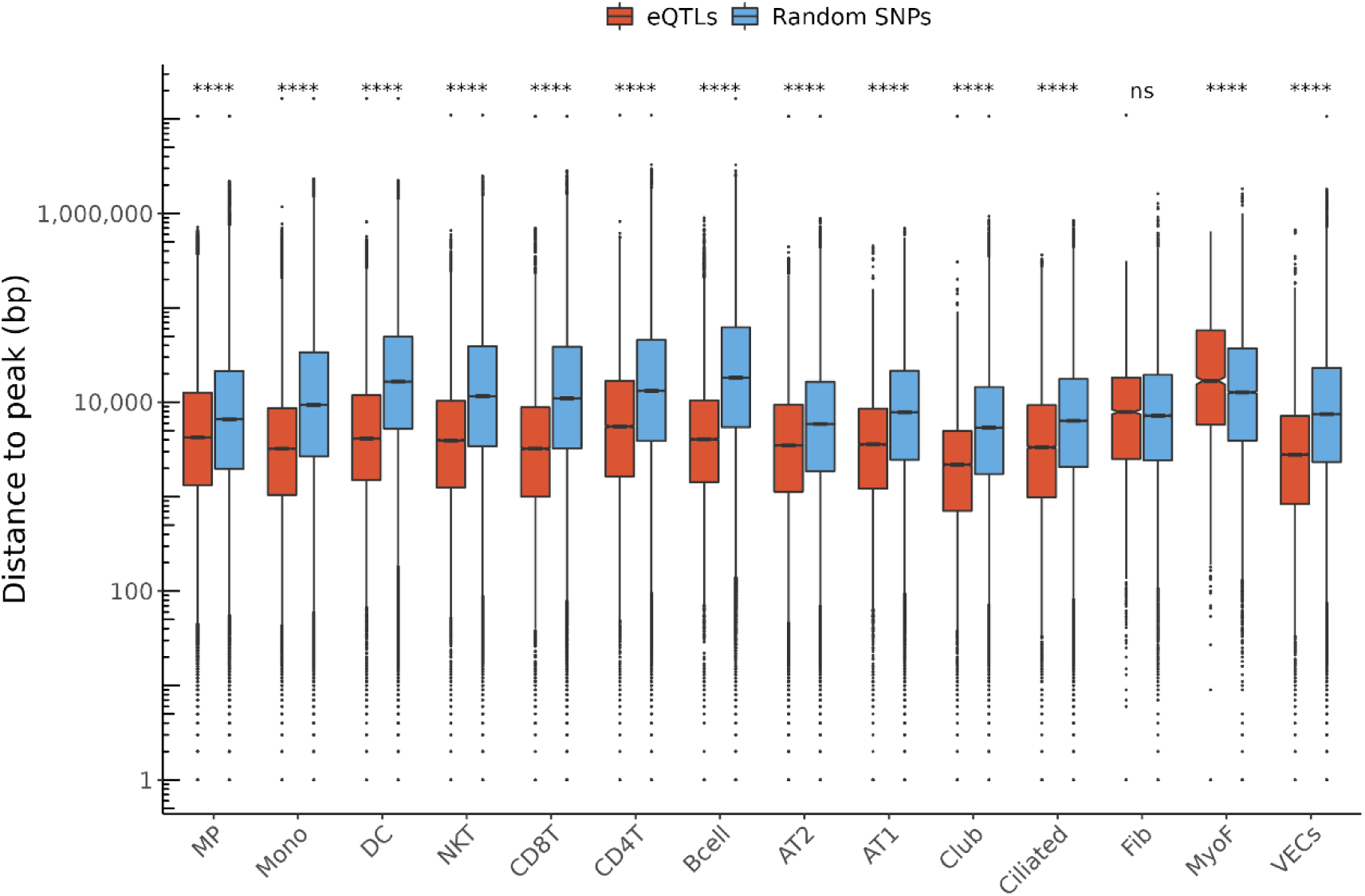
cis-eQTL distances to scATAC-seq peaks. Genomic distances from cis-eQTLs identified in each population to the nearest open chromatin region are shown in red. Random cis-SNPs tested for eQTL analysis are shown in blue. The y-axis in both plots shows the distances as log10 (base pairs + 1).

**Extended Data Figure 2.**
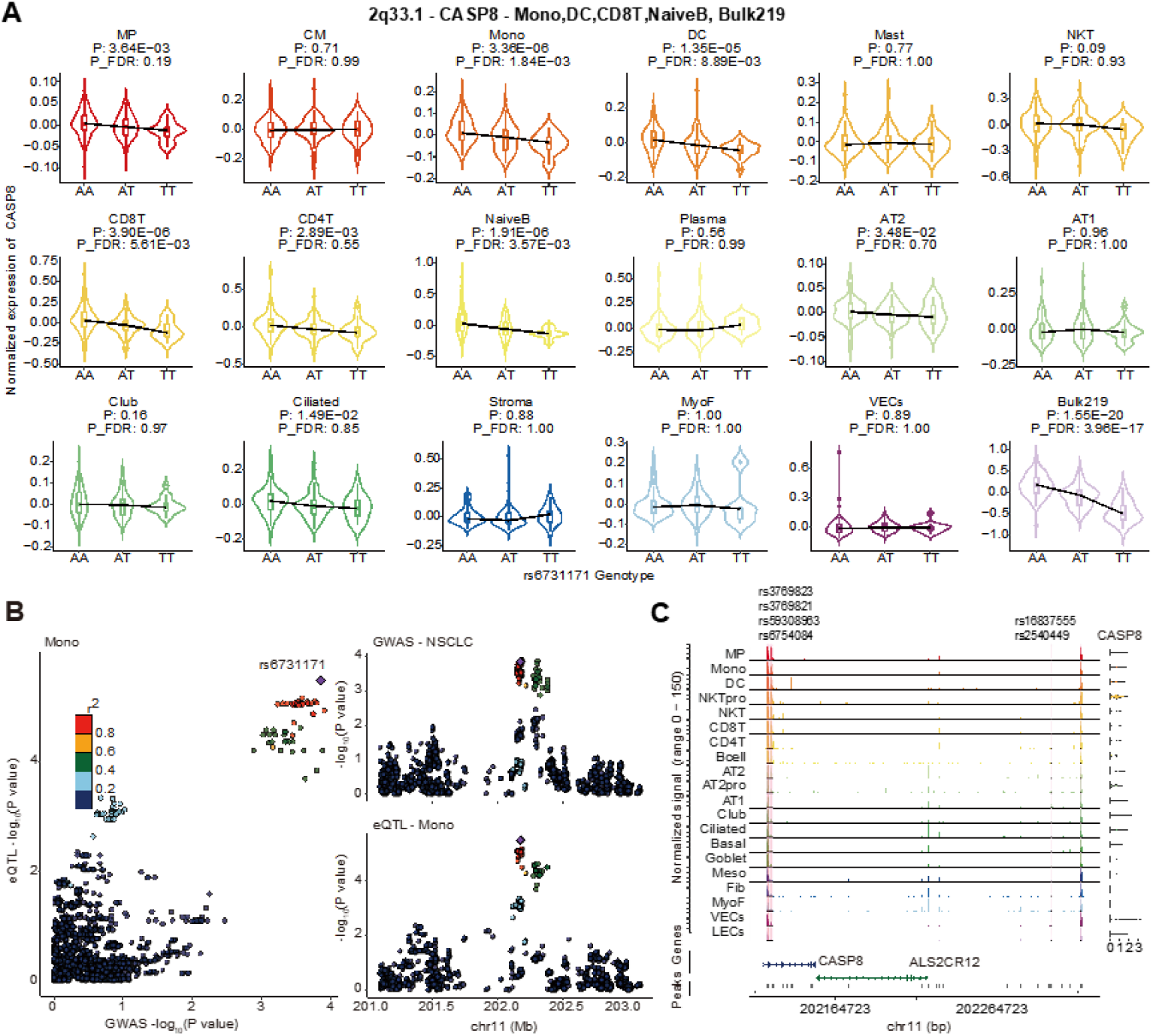
A detailed presentation of co-localization signals in the lung cancer GWAS loci resolved by immune cell population specificity. (A) Allelic violin diagrams show the effects of the lead variant rs6731171 regulating the expression of *CASP8* in specific cell types, such as Mono, DC, CD8T and NaiveB. (B) Regional plots show the overlap of GWAS and eQTL associations for *CASP8* in Mono. The lead variant is indicated with a purple diamond, and other points are colored according to the linkage disequilibrium index (r^2^ value) with the lead variant. (C) Sequencing tracks representing chromatin accessibility for the lead variants and their high LD variants (r² > 0.5), marked by rsIDs and vertical pink lines. Each track represents the aggregated snATAC signal of each cell type, normalized by the total number of reads in the regions. Arrow depicts the transcriptional direction of *CASP8*.

**Extended Data Figure 3.**
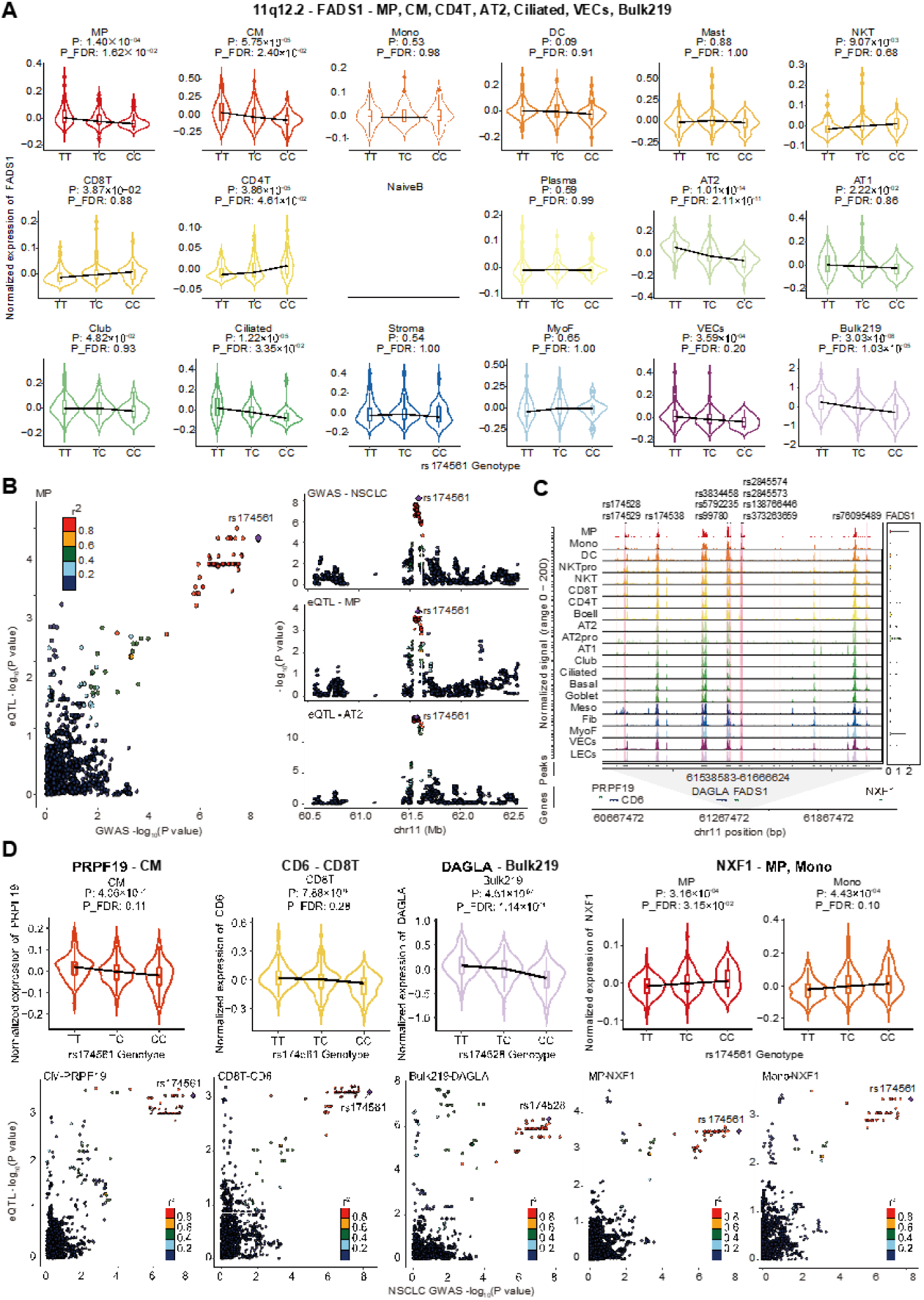
A detailed presentation of co-localization signals in the lung cancer GWAS loci resolved by broad cell population. (A) Allelic violin diagrams show the effects of the lead variant rs174561 regulating the expression of *FADS1* in specific cell types. (B) Regional plots show the overlap of GWAS and eQTL associations for *FADS1* in MP and AT2. The lead variant is indicated with a purple diamond, and other points are colored according to the linkage disequilibrium index (r^2^ value) with the lead variant. (C) Sequencing tracks representing chromatin accessibility for the lead variants and their high LD variants (r² > 0.5), marked by rsIDs and vertical pink lines. Each track represents the aggregated snATAC signal of each cell type, normalized by the total number of reads in the regions. Arrow depicts the transcriptional direction of *FADS1*. (D) Allelic violin diagrams show the effects of the lead variant regulating the expression of the target genes in specific cell types, such as *PRPF19*, *CD6*, *DAGLA* and *NXF1*. Regional plots show the overlap of GWAS and eQTL associations for those gene in corresponding subpopulations.

**Extended Data Figure 4.**
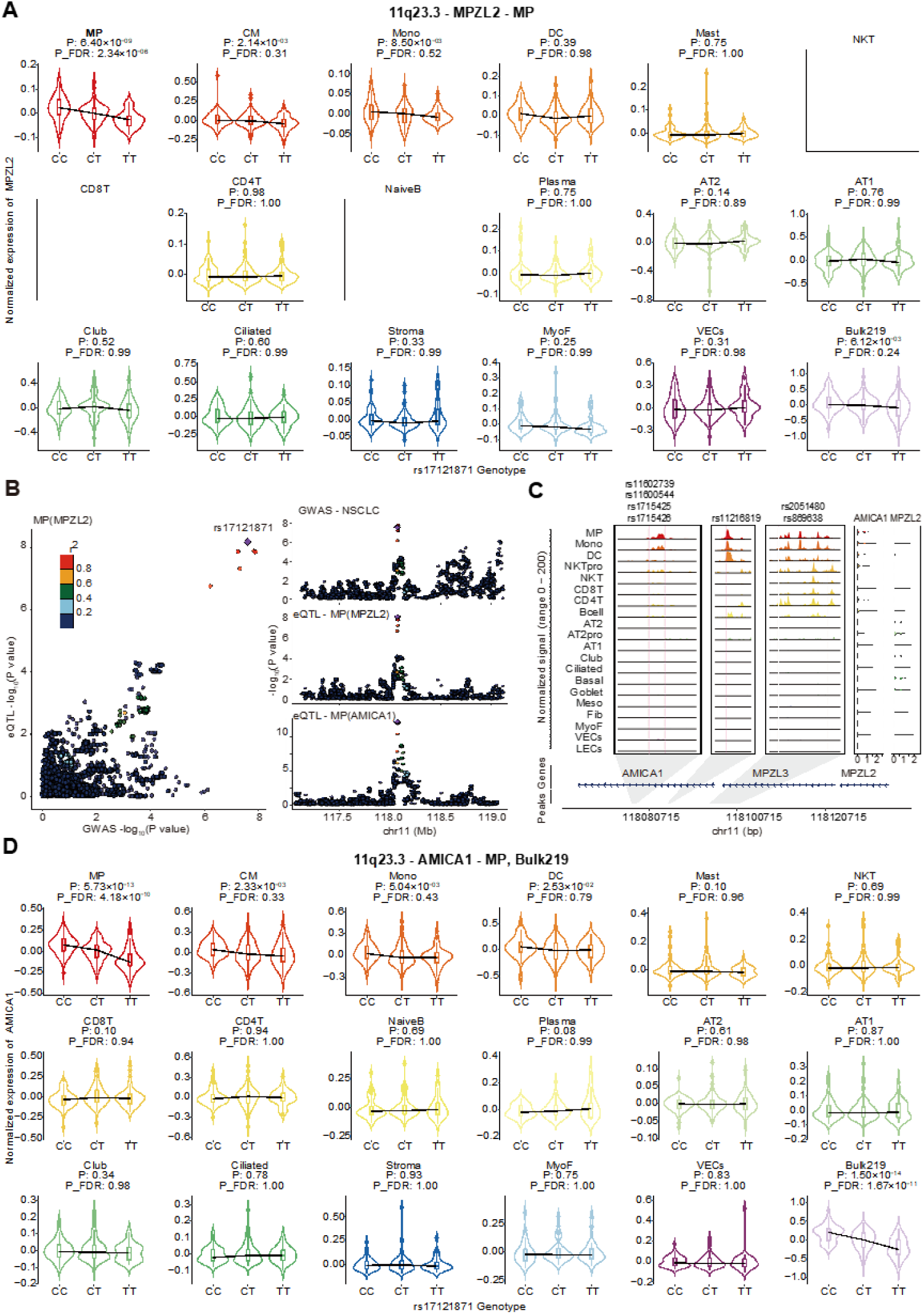
A detailed presentation of co-localization signals in the lung cancer GWAS loci resolved by MP cell specificity. (A) Allelic violin diagrams show the effects of the lead variant rs17121871 regulating the expression of *MPZL2* in specific cell types. (B) Regional plots show the overlap of GWAS and eQTL associations for *MPZL2* and *AMICA1* in MP. The lead variant is indicated with a purple diamond, and other points are colored according to the linkage disequilibrium index (r^2^ value) with the lead variant. (C) Sequencing tracks representing chromatin accessibility for the lead variants and their high LD variants (r² > 0.5), marked by rsIDs and vertical pink lines. Each track represents the aggregated snATAC signal of each cell type, normalized by the total number of reads in the regions. Arrow depicts the transcriptional direction of *AMICA1* and *MPZL2*. (D) Allelic violin diagrams show the effects of the lead variant rs17121871 regulating the expression of *AMICA1* in specific cell types.

**Extended Data Figure 5.**
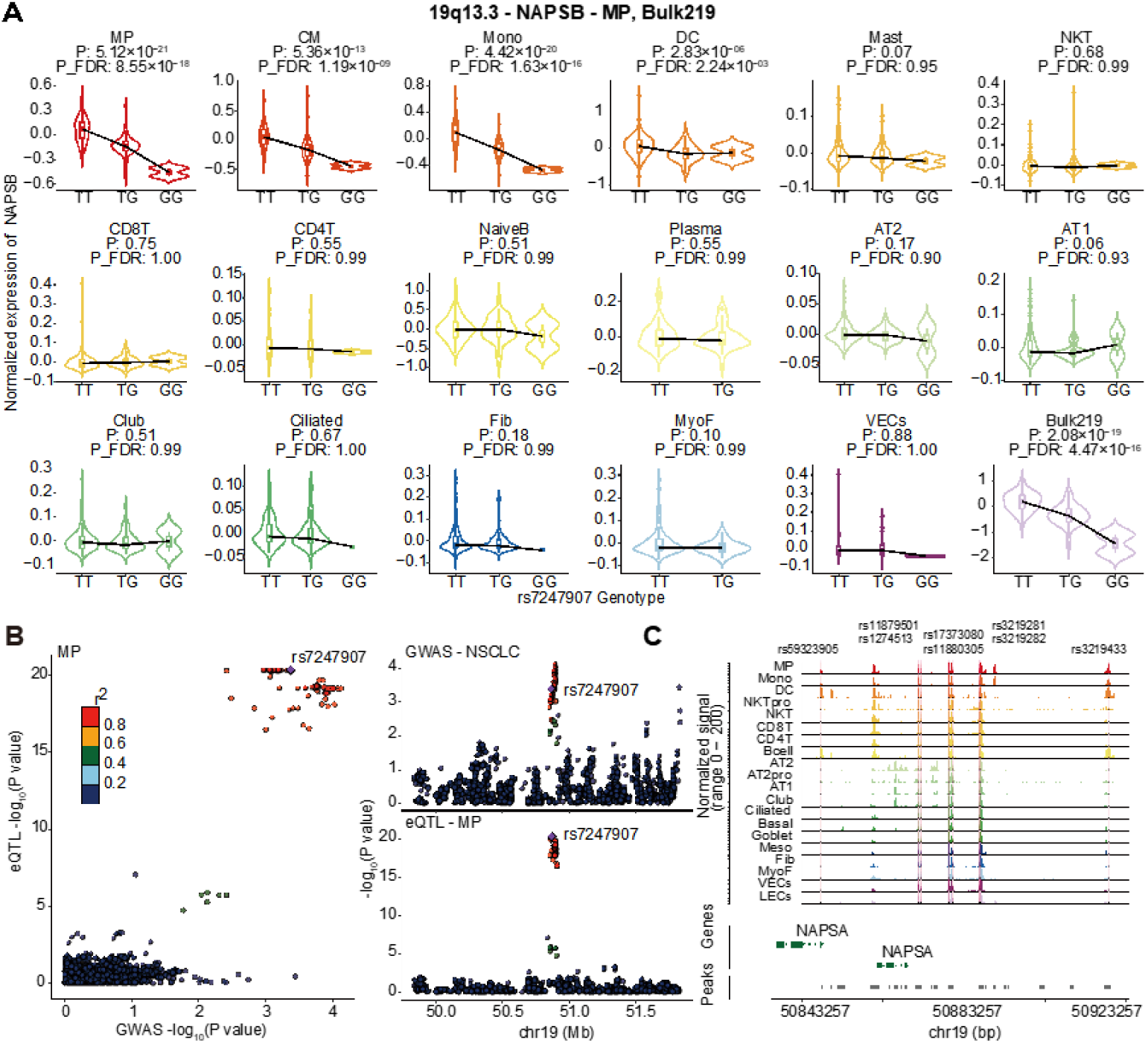
A detailed presentation of co-localization signals in novel lung cancer GWAS loci resolved by immune cell population specificity. (A) Allelic violin diagrams show the effects of the lead variant rs7247907 regulating the expression of *NAPSB* in specific cell types, such as Mono, DC, CD8T and NaiveB. (B) Regional plots show the overlap of GWAS and eQTL associations for *NAPSB* in MP. The lead variant is indicated with a purple diamond, and other points are colored according to the linkage disequilibrium index (r^2^ value) with the lead variant. (C) Sequencing tracks representing chromatin accessibility for the lead variants and their high LD variants (r² > 0.5), marked by rsIDs and vertical pink lines.. Each track represents the aggregated snATAC signal of each cell type, normalized by the total number of reads in the regions. Arrow depicts the transcriptional direction of *NAPSB*.

**Extended Data Figure 6.**
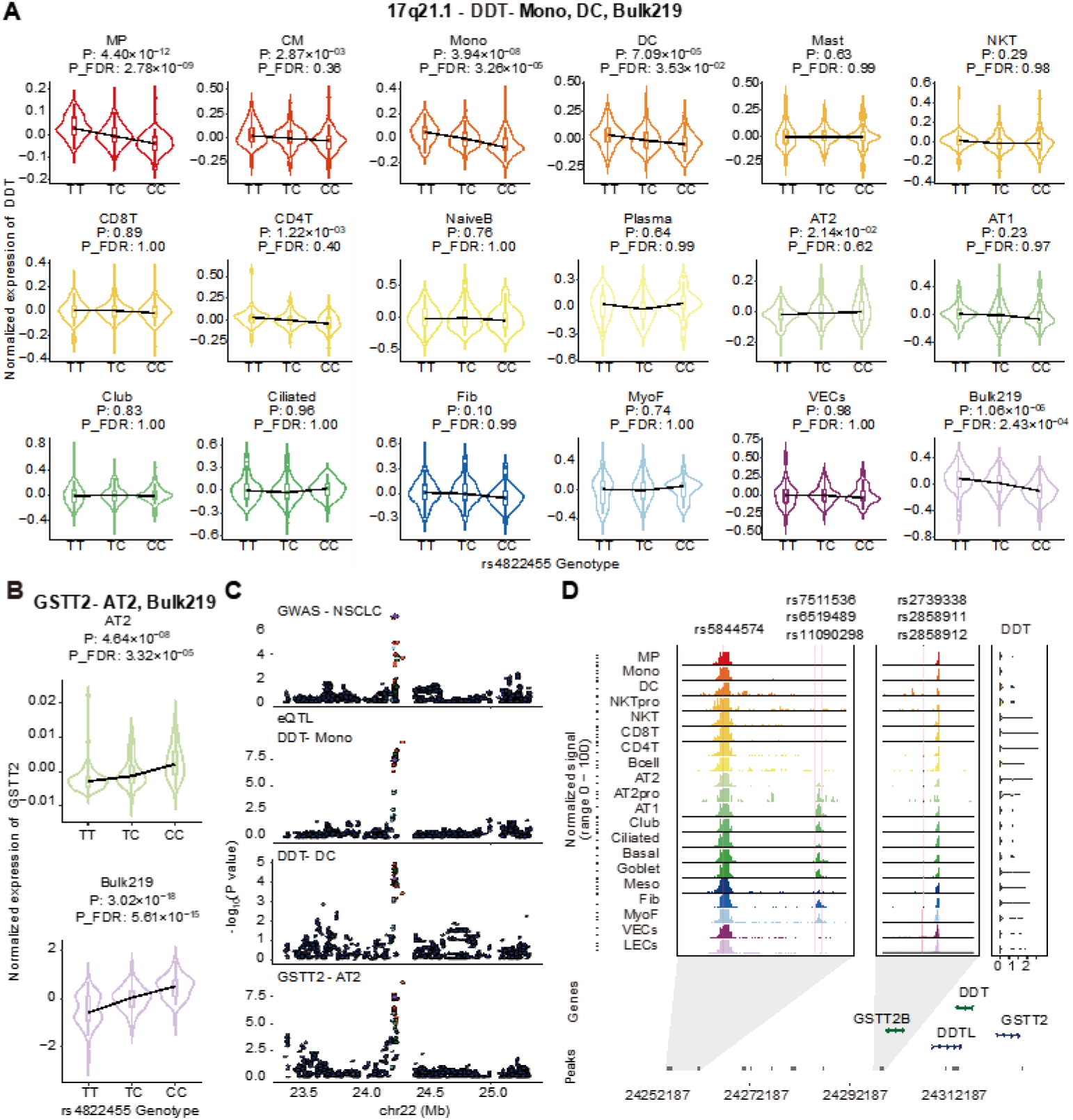
A detailed presentation of co-localization signals in novel lung cancer GWAS loci resolved by broad cell population. (A) Allelic violin diagrams show the effects of the lead variant rs4822455 regulating the expression of *DDT* in specific cell types. (B) Allelic violin diagrams show the effects of the lead variant rs4822455 regulating the expression of *GSTT2* in specific cell types. (C) Regional plots show the overlap of GWAS and eQTL associations for *DDT* in Mono and DC, *GSTT2* in AT2. The lead variant is indicated with a purple diamond, and other points are colored according to the linkage disequilibrium index (r^2^ value) with the lead variant. (D) Sequencing tracks representing chromatin accessibility for the lead variants and their high LD variants (r² > 0.5), marked by rsIDs and vertical pink lines. Each track represents the aggregated snATAC signal of each cell type, normalized by the total number of reads in the regions. Arrow depicts the transcriptional direction of *DDT* and *GSTT2*.

## Notes

### Competing Interest Statement

The authors have declared no competing interest.

